# CRK2 enhances salt tolerance by regulating callose deposition in connection with PLDα1

**DOI:** 10.1101/487009

**Authors:** Kerri Hunter, Sachie Kimura, Anne Rokka, Cuong Tran, Masatsugu Toyota, Jyrki P. Kukkonen, Michael Wrzaczek

## Abstract

High salinity has become an increasingly prevalent source of stress to which plants need to adapt. The receptor-like protein kinases (RLKs), including the cysteine-rich receptor-like kinase (CRK) subfamily, are a highly expanded family of transmembrane proteins in plants that are largely responsible for communication between cells and the extracellular environment. Various CRKs have been implicated in biotic and abiotic stress responses, however their functions on a cellular level remain largely uncharacterized. Here we have shown that CRK2 enhances salt tolerance at the germination stage in *Arabidopsis thaliana* and also modulates root length. We established that functional CRK2 is required for salt-induced callose deposition. In doing so, we revealed a novel role for callose deposition, in response to increased salinity, and demonstrated its importance for salt tolerance during germination. Using fluorescently tagged proteins we observed specific changes in CRK2’s subcellular localization in response to various stress treatments. Many of CRK2’s cellular functions were dependent on phospholipase D (PLD) activity, as were the subcellular localization changes. Thus we propose that CRK2 acts downstream of PLD during salt stress to promote callose deposition and regulate plasmodesmal permeability, and that CRK2 adopts specific stress-dependent subcellular localization patterns in order to carry out its functions.

**One sentence summary:** The receptor-like kinase CRK2 adopts PLDα1-dependent stress-induced subcellular localization patterns to regulate callose deposition at plasmodesmata, enhancing salt tolerance in *Arabidopsis thaliana*.

## Introduction

High soil salinity is becoming increasingly problematic in agriculture, with recent estimates assigning at least 20% of the total cultivatable land as affected (FAO and ITPS, 2015). In order to develop crops that can tolerate such conditions, it is first necessary to understand the mechanisms of salt stress responses and tolerance, much of which remains insufficiently characterized at the cellular and biochemical level. A high salinity environment exerts an osmotic stress on plants, as well as interfering with soil structure, nutrient and water acquisition, ionic balances, and solute concentrations within cells. This can lead to decreased plant growth, health, yield, and overall agricultural productivity (Machado and Serralheiro, 2017; Shrivastava and Kumar, 2015). Cellular responses to salt stress need to incorporate mechanisms to deal with the physical features of salt stress, such as membrane integrity and osmotic pressure, and the biochemical aspects, such as transport and balance of water, nutrients, solutes, and ions, while maintaining overall plant health and development. The multifaceted response of plant cells to high salinity is currently known to include: activation of NADPH oxidase respiratory burst homologs (RBOHs) and reactive oxygen species (ROS) production (Ma et al., 2012), calcium influx (Choi et al., 2014; Knight et al., 1997; Tracy et al., 2008), activation of phospholipase D (PLD) and phosphatidic acid (PA) production (Hong et al., 2010; Li et al., 2009), cell wall modifications (Tenhaken, 2015), changes in plasma membrane composition and formation of microdomains (Elkahoui et al., 2004; Hao et al., 2014; López-Pérez et al., 2009; Wu et al., 1998), and increased endocytosis of various receptors and channels (Baral et al., 2015), notably aquaporins to regulate water transport (Li et al., 2011; Luu et al., 2011; Ueda et al., 2016). The integration and regulation of these processes, however, still lacks a complete understanding.

The receptor-like protein kinases (RLKs) are a highly expanded family of transmembrane proteins in plants, and are largely responsible for the communication between cells and the extracellular environment. These proteins are widely represented across plant lineages, with *Arabidopsis thaliana* (Arabidopsis) containing more than 600 different RLKs (Shiu and Bleecker, 2003). The large diversity of RLKs and potential for crosstalk and interaction could permit responses to a huge variety of stimuli; accordingly, RLKs are known to regulate growth, development, and stress adaptation, including the response to pathogens and other biotic and abiotic stimuli (Kimura et al., 2017). RLKs are typically localized at the plasma membrane, with the N-terminal signal perception domain residing in the apoplast and the C-terminal kinase domain extending into the cytoplasm. This orientation permits sensing of extracellular ligands or microenvironment changes and subsequent transmission of the signal to the intracellular environment via the kinase activity or other protein interactions.

Cysteine-rich receptor-like kinases (CRKs) represent a subgroup of RLKs, consisting of 44 members in Arabidopsis (Wrzaczek et al., 2010). CRKs are defined by an extracellular domain containing two copies of the domain of unknown function 26 (DUF26) configuration of conserved cysteines C-X8-C-X2-C (Chen, 2001; Vaattovaara et al., 2019). Based on their expression profile (Wrzaczek et al., 2010) and loss-of-function phenotypes (Bourdais et al., 2015), CRKs are promising signalling candidates for both biotic and abiotic stress-responsive pathways. In particular, CRK2, 5, 8, 11, 28, 29, 36, 37, and 45 have been implicated in the response to salt stress (Bourdais et al., 2015; Tanaka et al., 2012; Zhang et al., 2013). While some CRKs have been linked to ROS signalling (Idänheimo et al., 2014) and cell death (Burdiak et al., 2015; Yadeta et al., 2017), for the majority the functions on a cellular and biochemical level remain largely uncharacterized.

In this study we sought to characterize the role of the receptor-like kinase CRK2 during salt stress. We show that CRK2 enhances germination and root length under conditions of high salinity. We describe how the protein is acting in connection with PLDα1 to regulate callose deposition in response to salt, and demonstrate that subcellular protein localization plays a major role in regulating CRK2 function. We also demonstrate novel salt-induced callose deposition, and its significance for salt tolerance.

## Results

### CRK2 interacts with proteins involved in salt responses

CRK2 was previously linked to multiple stress-related processes (Bourdais et al., 2015), however the mechanisms of its involvement on a biochemical and cellular level remained uncharacterized. RLKs typically do not act alone, but rather in protein complexes. Therefore, we performed a proteomics screen to identify proteins participating in CRK2-containing complexes as an initial step for further characterization of protein function. CRK2–YFP protein was immunoaffinity-purified from seedlings and interacting proteins were identified by mass spectrometry. Unspecific interactors were removed by comparison to the YFP–Myc control line to exclude proteins identified as interacting with YFP. The full list of identified proteins is available in Supplementary Table 1. Experiments were performed under standard growth conditions as well as NaCl and H_2_O_2_ treatments; however, in most cases no striking differences in identified proteins were noted between the different conditions. The majority of top interactors identified were plasma membrane associated proteins. Cytoplasmic proteins and extracellular proteins were also considered relevant, as CRK2 contains domains which extend into both the apoplast and cytoplasm. Nuclear or organelle localized proteins identified were considered probable contaminants, which likely came into contact with the bait CRK2-YFP during the protein extraction process.

**Table 1.**
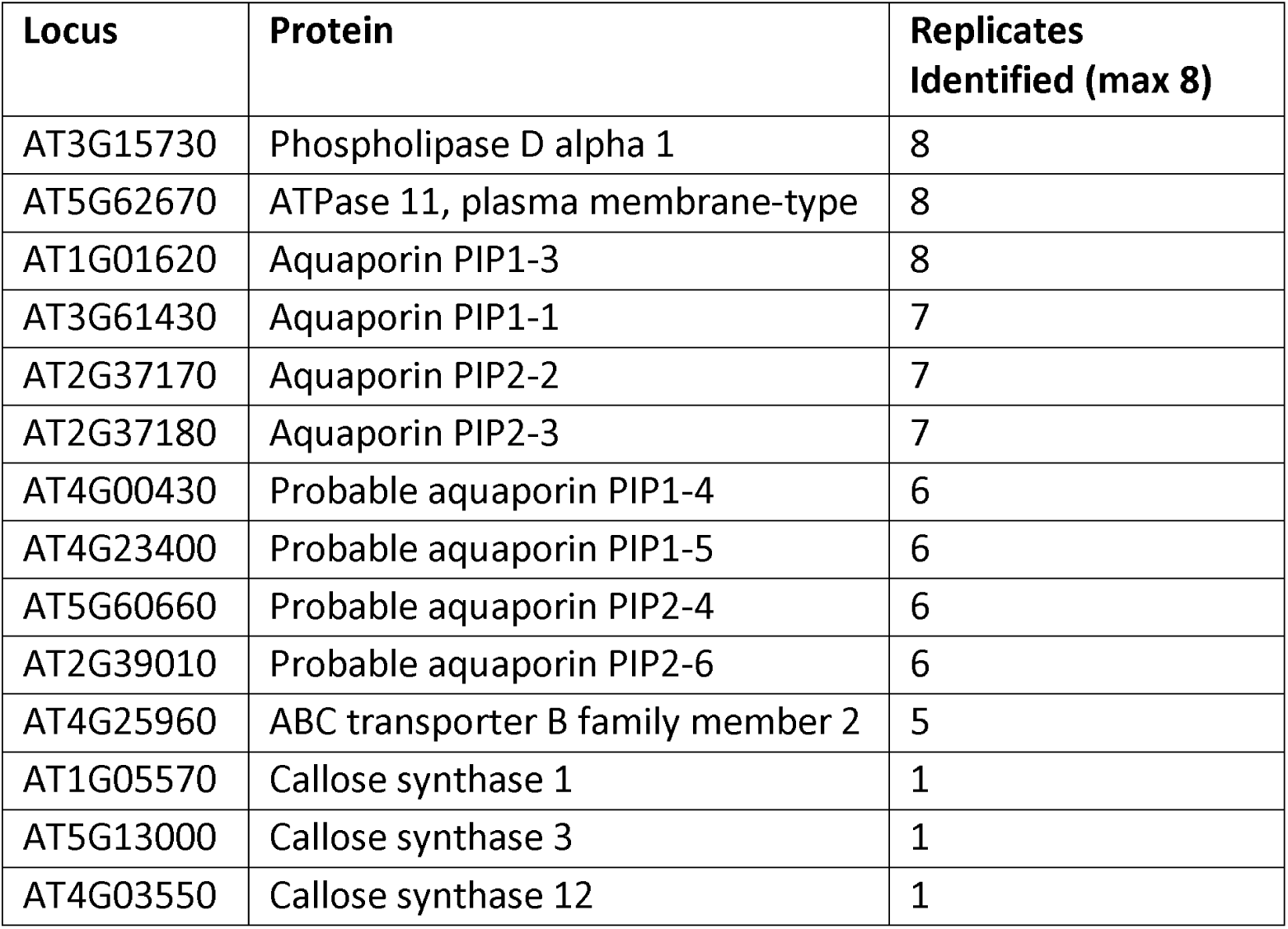
Selected proteins identified as interacting with CRK2. The 35S::CRK2–YFP_9-3 overexpression line was used in all replicates; unspecific interactors were removed by comparison to the 35S::YFP–Myc_9-6 control line. Each replicate indicates a separate immunoprecipitation in which the protein was identified.

| **Locus** | **Protein** | **Replicates Identified (max 8)** |
| --- | --- | --- |
| AT3G15730 | Phospholipase D alpha 1 | 8 |
| AT5G62670 | ATPase 11, plasma membrane-type | 8 |
| AT1G01620 | Aquaporin PIP1-3 | 8 |
| AT3G61430 | Aquaporin PIP1-1 | 7 |
| AT2G37170 | Aquaporin PIP2-2 | 7 |
| AT2G37180 | Aquaporin PIP2-3 | 7 |
| AT4G00430 | Probable aquaporin PIP1-4 | 6 |
| AT4G23400 | Probable aquaporin PIP1-5 | 6 |
| AT5G60660 | Probable aquaporin PIP2-4 | 6 |
| AT2G39010 | Probable aquaporin PIP2-6 | 6 |
| AT4G25960 | ABC transporter B family member 2 | 5 |
| AT1G05570 | Callose synthase 1 | 1 |
| AT5G13000 | Callose synthase 3 | 1 |
| AT4G03550 | Callose synthase 12 | 1 |

The identified proteins were annotated for gene ontology classifications based on biological process and analyzed by Singular Enrichment Analysis using the AgriGO GO Analysis Toolkit and Database (China Agricultural University; http://bioinfo.cau.edu.cn/agriGO/) to visualize the processes relevant to the CRK2 interactors (Supplementary Fig. S1, Supplementary Table 2). Several transmembrane channels or transport proteins were identified as top interactors, including multiple aquaporins, ABC type transporters, and ATPases (Supplementary Table 1), supporting a potential role for CRK2 in the mediation of cellular ionic or osmotic balances. Another protein family which was identified multiple times were the jacalin-related lectins (Supplementary Table 1), which have been shown to be involved in tolerance to both abiotic and biotic stresses (Esch and Schaffrath, 2017). It was shown that secreted proteins with a single DUF26 domain from Gingko and maize bind mannose *in vitro* (Ma et al., 2018; Miyakawa et al., 2014, 2009). In contrast, two related proteins containing tandem DUF26 domains, as is found in CRK2, were not able to bind mannose, and ligands of proteins with tandem DUF26 domains remain unknown (Vaattovaara et al., 2019). Therefore, it is conceivable that CRK2 and jacalin-related lectins might participate together in a complex for binding extracellular carbohydrates, glycopeptides, or other extracellular molecules.

**Table 2.** Chemicals and experimental conditions.

| **Chemical** | **Function** | **Concentration** | **Time** | **Source** |
| --- | --- | --- | --- | --- |
| 1-butanol | inhibits phosphatidic acid production by phospholipase D | 0.4% v/v | 10 min | Sigma-Aldrich, Darmstadt, Germany |
| 2-butanol | negative control for 1-butanol; does not affect phospholipase D | 0.4% v/v | 10 min | Alfa Aesar; ThermoFisher, Kandel, Germany |
| CaCl<sub>2</sub> | raises extracellular Ca concentration | 1 mM | 30 min | Merck, Darmstadt, Germany |
| Cyclopiazonic acid (CPA) | inhibits SERCA (sarcoendoplasmic reticulum calcium-ATPases), inducing calcium release and secondary store-operated Ca<sup>2+</sup> influx | 3 µM | 30 min | Tocris, Abingdon, United Kingdom |
| Dextromethorphan (DXM) | inhibits Ca<sup>2+</sup> and Na<sup>+</sup> channels | 10 µM | 10 min | RBI; Sigma-Aldrich, Darmstadt, Germany |
| Dimethyl sulfoxide (DMSO) | control for chemicals dissolved in DMSO | 1 µL | 30 min | Sigma-Aldrich, Darmstadt, Germany |
| Diphenyleneiodonium chloride (DPI) | inhibits flavoproteins (including RBOHs) | 10 µM | 1 h | Sigma-Aldrich, Darmstadt, Germany |
| Dyngo-4a | inhibits dynamin and clathrin-mediated endocytosis | 30 µM | 10 min | Abcam, Cambridge, United Kingdom |
| Ethylene glycol-bis(β-aminoethyl ether)-N,N,N',N'-tetraacetic acid (EGTA) | chelates extracellular Ca<sup>2+</sup> | 5 mM | 10 min | Sigma-Aldrich, Darmstadt, Germany |
| flg22 | mimics biotic stress | 10 µM | 30 min | GenScript, Piscataway, USA |
| H<sub>2</sub>O<sub>2</sub> | extracellular ROS treatment | 1 mM | 30 min | Sigma-Aldrich, Darmstadt, Germany |
| Ionomycin | Induces Ca<sup>2+</sup> influx | 10 µM | 30 min | Merck Millipore, Darmstadt, Germany |
| KCl | control for NaCl ionic component | 150 mM | 30 min | Fluka; Honeywell, Bucharest, Romania |
| LaCl<sub>3</sub> | inhibits Ca<sup>2+</sup> channels | 1 mM | 10 min | Fluka; Honeywell, Bucharest, Romania |
| Mannitol | osmotic stress, plasmolysis | 800 mM | 15 min | Alfa Aesar;<br>ThermoFisher, Kandel,<br>Germany |
| NaCl | salt stress | 150 mM | 30 min | Sigma-Aldrich,<br>Darmstadt, Germany |

Several interesting proteins were identified which were previously implicated in salt stress responses, and a list of selected interactors is presented in Table 1. The selected proteins were chosen based on high confidence scores from mass spectrometry identification, frequency of replicates identified in, and references to salt stress involvement in literature. Many of the top interactors have been linked to salt tolerance, including aquaporins (Bhardwaj et al., 2013), ATPases (Janicka-Russak and Kabała, 2015), and PHOSPHOLIPASE D ALPHA 1 (PLDα1; Bargmann et al., 2009). PLDα1 was identified as a CRK2-interacting protein in all eight replicates, and was consistently one of the top interactors (Table 1; Supplementary Table. 1). PLDα1 is mainly located in the cytoplasm in the resting state, and translocates to the plasma membrane upon activation (Wang et al., 2000; Zien et al., 2001). The C-terminal cytoplasmic domain of CRK2 could potentially mediate this docking, or alternatively, respond to PLDα1 docking. Three callose synthases were identified, and while callose deposition has not yet been explicitly documented during salt stress, it is a common feature to many other stress responses such as pathogen infection (Felix et al., 1999; Gómez-Gómez and Boller, 2000; Jacobs et al., 2003), heavy metal toxicity (O’Lexy et al., 2018), and osmotic stress (Xie et al., 2012). While the callose synthases were only identified in one replicate each, their large protein size and multiple transmembrane regions make these proteins inherently difficult to purify, and thus could account for the relatively low abundance in the samples.

### CRK2 enhances salt tolerance

Previous results indicated a role for CRK2 in salt stress responses, since the T-DNA insertion mutant *crk2* (Supplementary Fig. S2) exhibited decreased percentage of germination compared to Col-0 on media containing 150 mM NaCl (Bourdais et al., 2015). We confirmed that *crk2* is more salt-sensitive as assessed by percentage of germination (Fig. 1A), and demonstrated that this phenotype can be restored to wild type by complementation with CRK2–YFP expressed under its native promoter (pCRK2::CRK2–YFP_1-22 and 1-17, in *crk2* background; Fig. 1A). Overexpression of CRK2–YFP under the control of the CaMV 35S promoter (35S::CRK2–YFP_9-3, in Col-0 background) significantly increased salt tolerance at the germination stage (Fig. 1A).

**Fig. 1.**
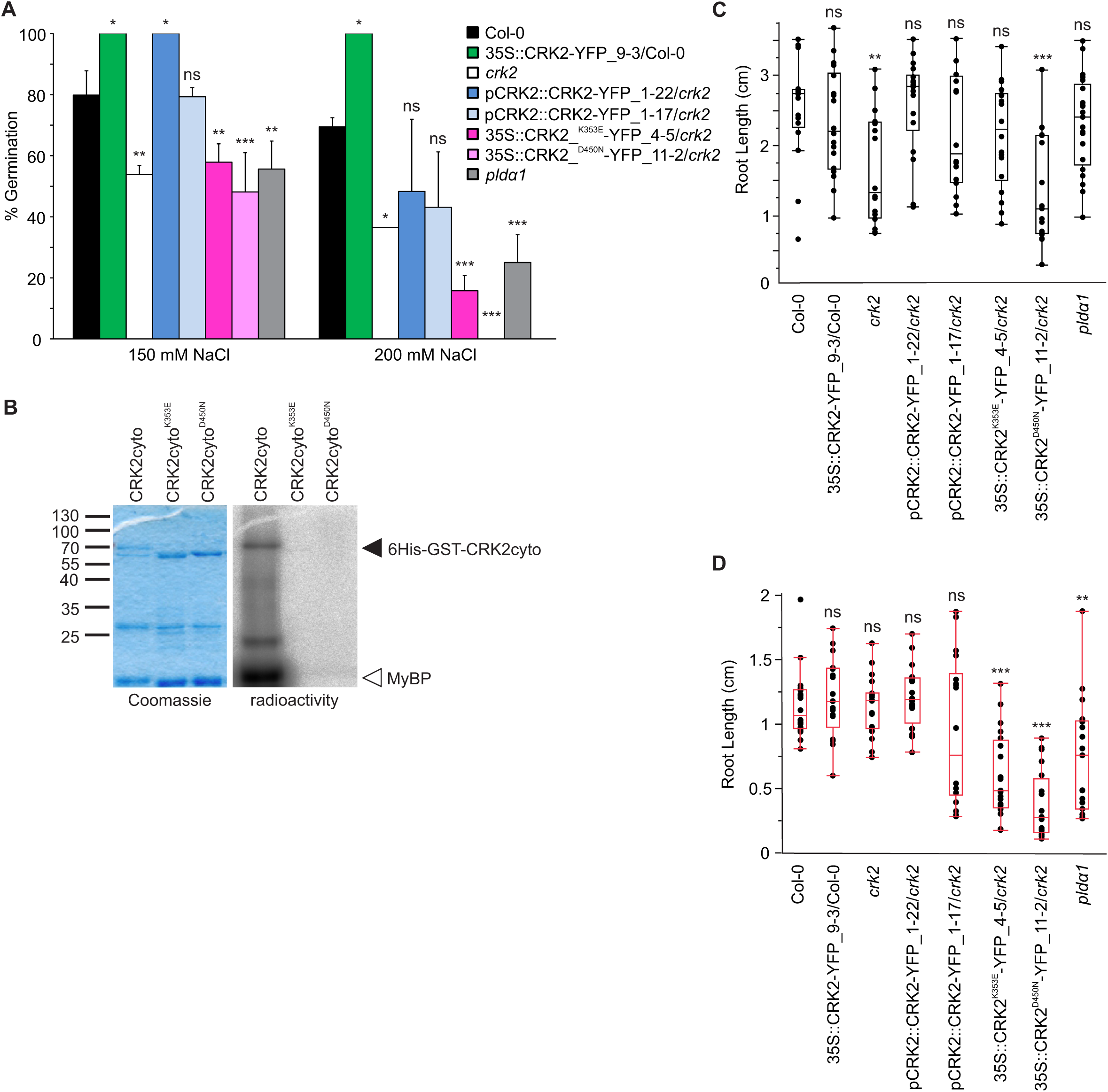
CRK2 enhances salt tolerance. (A) Overexpression of CRK2 increases salt tolerance at the germination stage, loss of functional CRK2 reduces salt tolerance. Data was normalized to the untreated controls for each line. Comparisons are to Col-0 (one-way ANOVA, *post hoc* Dunnett); n = 3; error bars indicate standard deviation. (B) CRK2 is an active kinase *in vitro*; kinase-dead protein variants lack kinase activity. (C-D) CRK2 is involved in primary root elongation under standard growth conditions (C) and 150 mM NaCl (D). Comparisons are to Col-0 (one-way ANOVA, *post hoc* Dunnett); eight-day-old seedlings, transplanted to treatments at five days; n = at least 16. ns not significant, * P < 0.05, ** P < 0.01, *** P < 0.001.

CRK2 contains the conserved motifs of a typical kinase domain (Kornev et al., 2006; Stone and Walker, 1995). Using the soluble cytosolic region of CRK2 (CRK2cyto), tagged with GST, we demonstrated that CRK2 is an active kinase *in vitro*, and is capable of autophosphorylation as well as phosphorylation of the generic kinase substrate myelin basic protein (MyBP; Fig 1B). The two mutated variants of CRK2 (CRK2cyto^K353E^ and CRK2cyto^D450N^), which were designed to be kinase-dead, did not exhibit kinase activity *in vitro* (Fig 1B). These kinase-dead point mutations disable two different motifs typically required for an active kinase: the K353E mutation disrupts the ATP-binding site, whereas the D450N mutation disrupts the catalytic core. Western blot analysis confirmed expression of CRK2–YFP in all transgenic lines (Supplementary Fig. S3A). In order to compare relative protein amounts, the mean intensity of western blot bands was quantified and normalized to RUBISCO (ribulose-1,5-bisphosphate carboxylase/oxygenase) and Histone H3 as internal controls (Supplementary Fig. S3B).

The germination response to salt is dependent on CRK2 kinase activity; expression of mutated CRK2 variants (kinase-dead: 35S::CRK2^K353E^–YFP and 35S::CRK2^D450N^–YFP, in *crk2* background) failed to restore the wild type germination phenotype. In fact, the kinase-dead lines displayed even more severe salt-sensitivity than *crk2* (Fig. 1A). The higher salt concentration of 200 mM magnified the differences between lines, although the overall trend remained largely the same at both concentrations (Fig. 1A). Since PLDα1 was identified as a top interactor for CRK2 we also investigated its role in salt stress. The *pldα1* mutant line (Supplementary Fig. S2) has been previously characterized as salt-sensitive and defective in several cellular processes related to the salt stress response (Bargmann et al., 2009; Hong et al., 2016; Yu et al., 2010; Zhang et al., 2012). Here we show that *pldα1* has decreased germination on NaCl-containing media, with a similar phenotype as the *crk2* and CRK2 kinase-dead lines (Fig. 1A).

In addition to germination rate, changes in root length and morphology are also associated with salt stress (Bayazid et al., 2016; Julkowska et al., 2014; Kawa et al., 2016). Assessment of primary root length revealed differences between the CRK2 lines, when grown on both untreated and salt-containing media (Fig. 1C and D). The *crk2* and CRK2^D450N^ lines had significantly shorter roots under standard growth conditions compared to Col-0 (Fig. 1C). Under high salt conditions, both CRK2 kinase-dead lines had significantly shorter roots compared to Col-0 (Fig. 1D). The shorter root phenotype was complemented by expression of CRK2–YFP under its native promoter (Fig. 1C and D). Overexpression of CRK2–YFP under the 35S promoter also complemented the mutant phenotype, but did not further increase root length over that of wild type or native CRK2 expression (Fig. 1C and D). The *pldα1* mutant displayed decreased root length compared to Col-0 following NaCl treatment (Fig. 1D). Thus, CRK2 and PLDα1 appear to also be involved in the root length aspect of salt tolerance, and our results suggest that CRK2 kinase activity is important for this function.

NaCl treatment exerts both an osmotic and ionic stress on cells. In order to determine which of these components was more important in relation to CRK2, we tested germination on media containing mannitol or KCl. The results with mannitol were similar to NaCl, whereby overexpression of CRK2 leads to higher tolerance (Supplementary Fig. S4). However, *crk2* did not significantly differ from Col-0 when germinated on mannitol (Supplementary Fig. S4). Germination with KCl did not produce any significant differences between the three lines (Supplementary Fig. S4). This suggests that both the osmotic component and Na^+^ ionic toxicity contribute to the CRK2-mediated NaCl stress response.

### CRK2 protein re-localizes in response to stress, to distinct spots resembling plasmodesmata following NaCl treatment

CRK2 is a transmembrane protein and, like other RLKs, was predicted to localize to the plasma membrane based on the presence of an N-terminal localization signal sequence (Shiu and Bleecker, 2003). Subcellular protein localization was evaluated by live cell imaging using plants expressing a 35S::CRK2–YFP fusion protein. Under control conditions CRK2 localized to the cell periphery in epidermal cells (Fig. 2A), in contrast to YFP alone which localized to the cell periphery, cytoplasm, and nucleus (Fig. 2A). Plasmolysis of cells showed the presence of Hechtian strands (arrows, Fig. 2A), strongly supporting plasma membrane localization of CRK2–YFP. The CRK2 kinase-dead variants displayed a subcellular localization at the plasma membrane similar to that of the wild type protein (Supplementary Fig. S5).

**Fig. 2.**
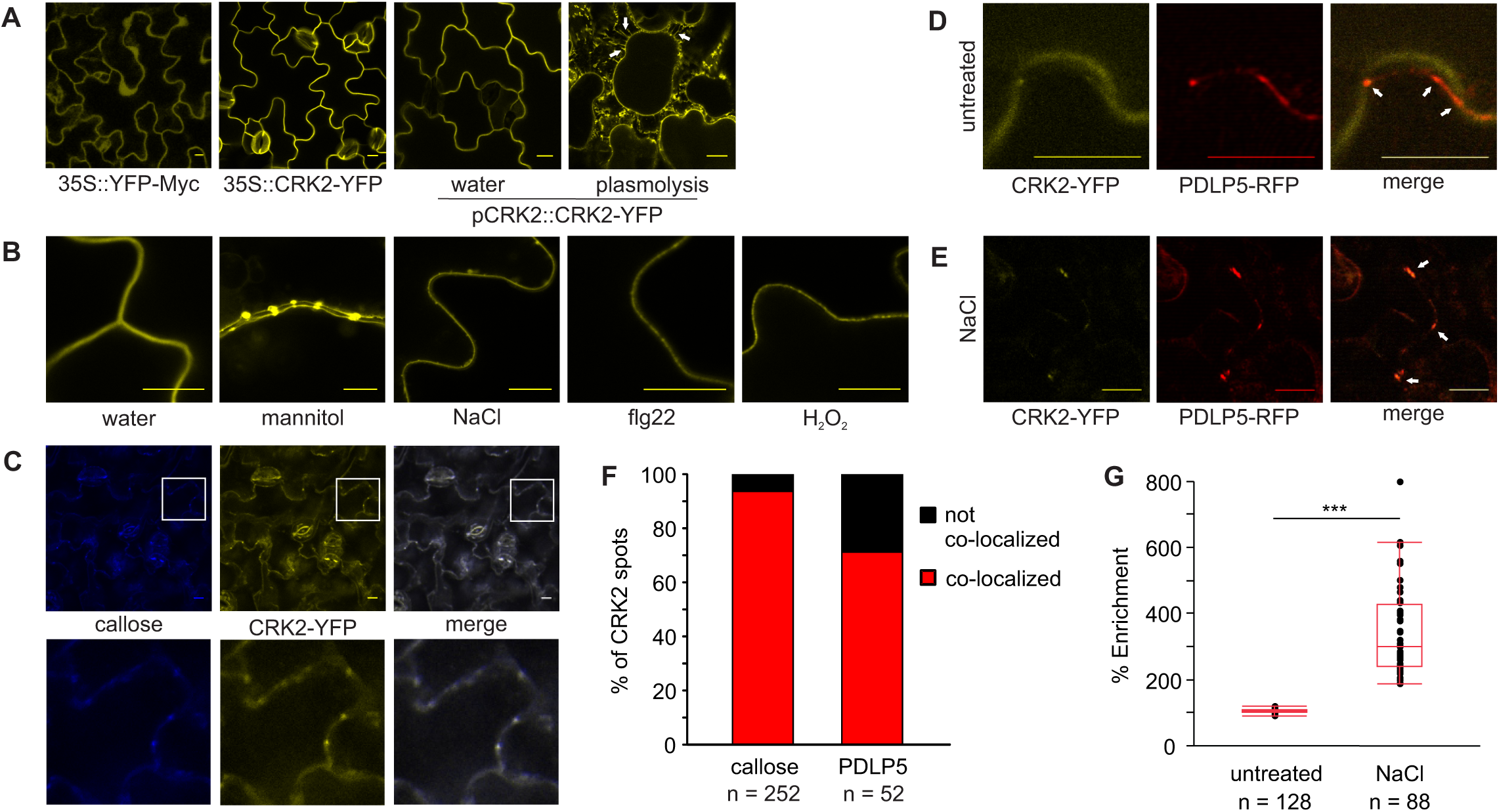
CRK2 subcellular protein localization. (A) CRK2-YFP localizes uniformly to the plasma membrane under standard growth conditions. Arrows indicate presence of Hechtian strands following plasmolysis. (B) 35S::CRK2-YFP re-localizes in response to abiotic and biotic stresses, to distinct stress-specific patterns along the plasma membrane. (C) Co-localization with callose deposits supports NaCl-induced plasmodesmal localization of CRK2-YFP; white box indicates zoomed area. (D) CRK2-YFP does not co-localize (arrows) with plasmodesmal marker PDLP5-RFP under standard growth conditions. (E) CRK2-YFP partially co-localizes (arrows) with PDLP5-RFP following NaCl treatment. (F) Quantification of CRK2-YFP co-localization with callose deposits and PDLP5-RFP following NaCl treatment. (G) Quantification of NaCl-induced re-localization of CRK2-YFP by percent enrichment at re-localization domains; *** P < 0.001 (one-way ANOVA, pooled t-test). Seven-day-old seedlings, epidermal cells; mannitol 800 mM 15 min, NaCl 150 mM 30 min, flg22 10 µM 30 min, H_2_O_2_ 1 mM 30 min. Scale bar = 10 µm.

Controlling protein localization within specific cellular compartments or domains is one means by which cells can regulate protein function post-translationally, and adjust in response to a stimulus. Localization to specialized domains along the plasma membrane has been observed for other RLKs, including FLAGELLIN SENSITIVE 2 (FLS2) and BRASSINOSTEROID INSENSITIVE 1 (BRI1; Bücherl et al., 2017). The subcellular localization of CRK2 changed in response to both abiotic and biotic stimuli. The protein assumed a new localization in spots along the plasma membrane, the size and pattern of which depended on the nature of the stress treatment (Fig. 2B). CRK2 expressed under its native promoter showed the same patterns as under the 35S promoter (Supplementary Fig. S6; compare to Fig. 2B), therefore the overexpression line was used for all further analysis of localization. Following treatment with mannitol or NaCl, CRK2 adopted a pattern of concentrated spots along the plasma membrane reminiscent of plasmodesmal localization (Fig. 2B) (Thomas et al., 2008; Lee et al., 2011; Xu et al., 2017; Diao et al., 2018). Much of the work on re-localization of RLKs has been carried out with microbe-associated molecular patterns (MAMP) treatments. Treatment with flg22, to mimic biotic stress, or H_2_O_2_, to raise the extracellular ROS concentration, produced a localization pattern of smaller, more frequent spots, possibly representing some form of microdomains (Fig. 2B). Localization at plasmodesmata following NaCl treatment was confirmed by co-localization of CRK2–YFP with callose deposits (Fig. 2C), which is often used as a plasmodesmata marker (Gaudioso-Pedraza & Benitez-Alfonso, 2014; Widana Gamage & Dietzgen, 2017; Xu et al., 2017). Quantification of co-localization revealed that 93.7% of CRK2 spots co-localized with callose deposits following NaCl treatment (Fig. 2F). CRK2–YFP also showed partial co-localization with PLASMODESMATA-LOCATED PROTEIN 5 (PDLP5), which has been previously identified as having a plasmodesmal localization (Lee et al., 2011). CRK2–YFP and PDLP5–RFP did not co-localize under standard growth conditions (Fig. 2D), however, co-localization increased following NaCl treatment (Fig. 2E). The co-localization of CRK2 spots with PDLP5 was 71.2% following NaCl treatment (Fig. 2F). Quantification of CRK2–YFP re-localization was achieved by calculating the percent enrichment at the re-localization domains compared to the rest of the plasma membrane. In untreated samples there were no discernable differences among different plasma membrane regions, leading to a 1:1 plasmodesmata:non-specific plasma membrane distribution (Fig. 2G). The enrichment of CRK2–YFP at discernable plasma membrane domains increased significantly following NaCl treatment, with a mean value of 3.4-fold enrichment at plasmodesmata (Fig. 2G).

### CRK2 re-localization is dependent on intracellular Ca^2+^ and PLD activity

To study the mechanism of CRK2’s stress-induced re-localization patterns, we first investigated the need for kinase activity using a kinase-dead variant of CRK2. No changes in localization were observed following NaCl, flg22, or H_2_O_2_ treatment of the kinase-dead line (Fig. 3A), establishing that while CRK2 kinase activity is not required for its delivery to the plasma membrane it requires an active kinase domain for the re-localization to occur.

**Fig. 3.**
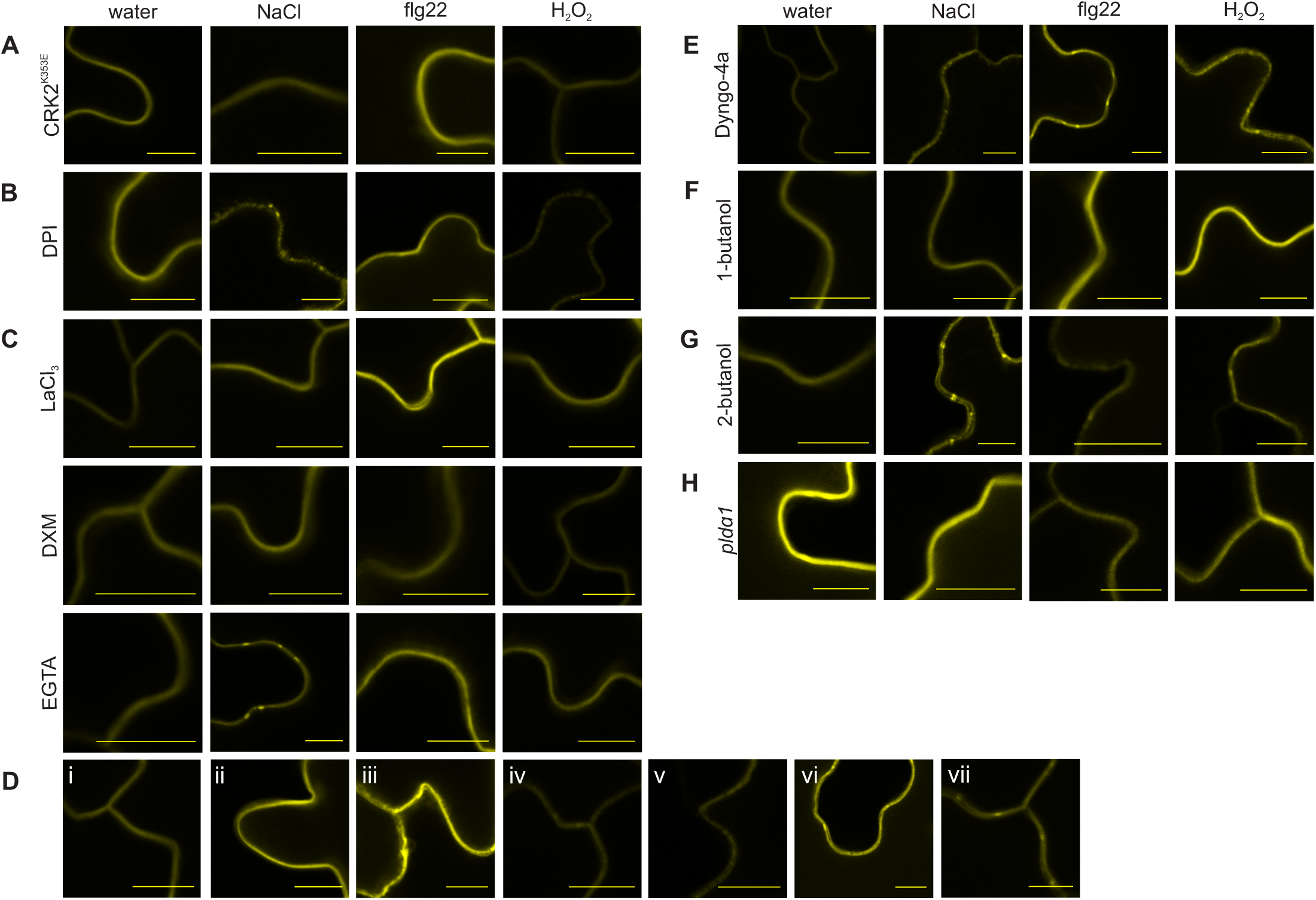
Mechanism of CRK2 stress-dependent localization changes. (A) Kinase activity is required for both abiotic and biotic stress-induced re-localization. (B) NADPH dependent ROS production is required for the biotic response, but not the abiotic re-localization. (C) Increased cytosolic calcium is required for both abiotic and biotic re-localization. (D) Increased cytosolic calcium is sufficient to induce CRK2 re-localization; (i) DMSO control, (ii) CaCl_2_, (iii) CaCl_2_ + ionomycin, (iv) CPA, (v) DPI + CaCl_2_ + ionomycin, (vi) DPI + CPA, (vii) CRK2^K353E^ + CaCl_2_ + ionomycin. (E) Clathrin-mediated internalization is not required for either abiotic or biotic re-localization. (F-G) PLD activity is required for both abiotic and biotic re-localization. (H) PLDa1 is required for both abiotic and biotic re-localization. 35S overexpression lines used in all replicates; seven-day-old seedlings, epidermal cells; treatment times and conditions according to Table 2. Scale bar = 10 µm.

ROS and Ca^2+^ are rapidly induced messengers common to numerous stress responses, and they couple to various downstream cellular events. Therefore, we investigated the influence of these components on CRK2 localization using an inhibitor-based approach (Table 2) whereby the samples were first pre-treated with the inhibitor, then subjected to the stress treatments and assessed for localization changes. Inhibition of extracellular ROS production by RBOHs was achieved with diphenyleneiodonium chloride (DPI), which inhibits flavoenzymes. Under these conditions CRK2–YFP was still able to re-localize upon NaCl treatment, but not upon flg22 treatment (Fig. 3B). This reveals a distinction between the abiotic and biotic stress responses not only in the pattern of localization, but also the mechanism. Treatment with H_2_O_2_ following DPI pre-treatment could still induce the spotted localization response (Fig. 3B). Reduction of calcium signalling, by blocking Ca^2+^ channels with LaCl_3_ or dextromethorphan (DXM), abolished the re-localization response of CRK2–YFP upon both biotic and abiotic stress (Fig. 3C). Chelating extracellular Ca^2+^ with EGTA had a similar effect, however some re-localization was still observed upon NaCl treatment. One explanation is that the Ca^2+^ channel blockers are more efficient than EGTA at preventing Ca^2+^ influx. Alternatively, this could indicate that intracellular Ca^2+^ release also plays a role in addition to Ca^2+^ influx from the extracellular environment (Fig. 3C). Treatment with H_2_O_2_ could no longer induce the response after inhibition of Ca^2+^ channels or chelation of extracellular Ca^2+^, suggesting a presiding need for calcium over ROS for the localization of CRK2–YFP (Fig. 3C). The need for a cytosolic Ca^2+^ increase was further supported by the observation that extracellular 1 mM CaCl_2_ alone was not enough to trigger the re-localization. (Fig. 3D-ii). Cytosolic Ca^2+^ elevation from the apoplast side was achieved by adding CaCl_2_ + ionomycin and from the intracellular stores by cyclopiazonic acid (CPA); in both cases re-localization was triggered (Fig. 3D-iii and -iv, respectively). Furthermore, Ca^2+^ elevation was stimulated while under the influence of DPI, to block NADPH oxidase-dependent ROS production, and again elevated cytosolic Ca^2+^ was enough to cause re-localization (Fig. 3D-v and -vi, respectively). Ca^2+^ elevation also restored the re-localization response in the kinase-dead line (Fig. 3D-vii). These results suggest that elevated intracellular Ca^2+^ is required and sufficient to induce the re-localization of CRK2, and likely serves as the primary signal for stress-induced CRK2 localization.

Next we investigated whether endocytosis was required for CRK2 re-localization, as many RLKs internalize as part of their regulation or signalling functions (Geldner and Robatzek, 2008). Dyngo-4a acts as a dynamin inhibitor to inhibit clathrin-mediated endocytosis (McCluskey et al., 2013). We first tested its effectiveness using the FLS2 receptor, for which internalization upon binding its ligand flg22 is well characterized (Robatzek et al., 2006). Dyngo-4a successfully prevented FLS2–GFP internalization following flg22 treatment and thus functions well in plant cells (Supplementary Fig. S7). Dyngo-4a treatment did not inhibit CRK2–YFP re-localization in response to NaCl, flg22, or H_2_O_2_, suggesting that clathrin-mediated endocytosis is not required for this process (Fig. 3E).

Finally, we examined the involvement of PLD, as these enzymes are capable of altering membrane composition and therefore potentially affecting the localization of plasma membrane proteins. We tested the requirement of PLD activity using 1-butanol as an inhibitor of PLD-based phosphatidic acid (PA) production. Primary alcohols such as 1-butanol inhibit PLD signalling by acting as a substrate 100-fold preferred over water for utilization in PLD hydrolysis, forcing the reaction in favor of producing phosphatidylbutanol instead of PA (Gardiner et al., 2003; Morris et al., 1997). Pre-treatment with 1-butanol effectively blocked CRK2–YFP re-localization in response to NaCl, flg22, and H_2_O_2_, establishing the requirement of PLD-based PA production for CRK2’s localization response during both abiotic and biotic stress (Fig. 3F). Secondary alcohols, such as 2-butanol, do not affect PLD activity, thus 2-butanol was used as a negative control. As expected, CRK2–YFP re-localization in response to NaCl, flg22, and H_2_O_2_ was not affected by pre-treatment with 2-butanol (Fig. 3G). CRK2–YFP transiently expressed in Col-0 seedlings exhibited re-localization responses comparable to the stable expression line (Supplementary Fig. S8), demonstrating that the transient expression system does not hinder CRK2 re-localization. CRK2–YFP transiently expressed in the *pldα1* mutant background was not able to re-localize following NaCl, flg22, or H_2_O_2_ treatments (Fig. 3H). Thus, PLDα1 is likely the major PLD isoform facilitating the CRK2 re-localization response to stress treatments.

### CRK2 is required for salt-induced callose deposition

The plasmodesmal localization of CRK2 following salt treatment and the identification of callose synthases as interacting partners prompted the investigation of CRK2’s effect on callose deposition. Callose deposition is commonly studied in the context of a stress response, and changes in callose profiles have been observed following bacterial and fungal infection, as well as osmotic stress (Felix et al., 1999; Gómez-Gómez and Boller, 2000; Jacobs et al., 2003; Xie et al., 2012). However, the callose deposition in response to acute salt stress has not yet been characterized. We could show that in wild-type Col-0 plants there was a significant increase in callose deposition in response to NaCl (Fig. 4A and B). This response was exaggerated in plants overexpressing CRK2 and lacking in the *crk2* and kinase-dead lines, suggesting that functionally active CRK2 protein is required for a salt-induced callose response (Fig. 4A and B). We investigated the importance of callose deposition for salt tolerance by assessing germination of the *cals1* mutant (*cals1-5*; Supplementary Fig. S2), which lacks functional CALLOSE SYNTHASE 1 (CALS1). CALS1 is involved in stress response, and regulation of plasmodesmata permeability by CALS1 following pathogen infection and mechanical wounding was demonstrated by (Cui and Lee, 2016). CALS1 is also one of the callose synthases found to interact with CRK2 and was used here as a representative, since no suitable mutant lines were available for the other identified callose synthases. Germination of *cals1* was reduced on media containing NaCl compared to Col-0 (Fig. 4C). This trend was observed at both concentrations of NaCl, however the difference was only statistically significant at the higher 200 mM concentration (Fig. 4C). The germination defect was not as severe as with *crk2* (compare to Fig. 1A), likely because of redundancy from the other callose synthases still present. The germination response of additional *cals1-2* and *cals1-3* alleles was similar to *cals1-5* and is shown in Supplementary Fig. S9. Salt-induced callose deposition is also lacking in *cals1* (Fig. 4D). Together, these results further highlight a role for callose deposition during salt stress, and support CALS1 as a major contributor to salt-induced callose deposition.

**Fig. 4.**
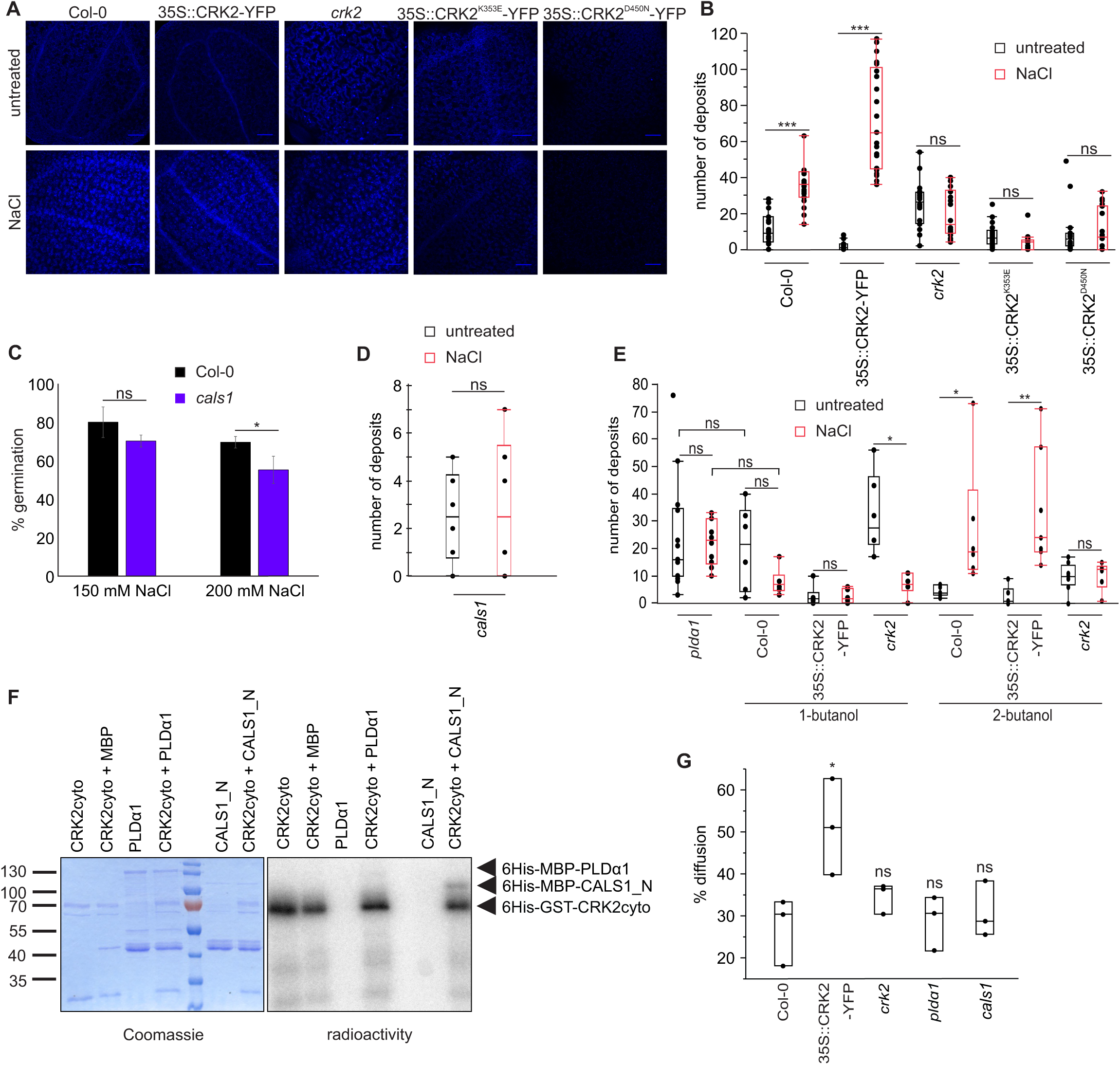
CRK2 is required for salt-induced callose deposition. (A) Aniline blue staining for callose deposition. Scale bar = 100 µm. (B) Kinase-active CRK2 is required for NaCl-induced callose deposition. Quantification of callose deposits; comparisons are between untreated and NaCl-treated samples for each line (one-way ANOVA, *post hoc* Tukey HSD); n = at least 15. (C) Germination of cals1 mutant is reduced on salt-containing media. Comparisons are to Col-0 (one-way ANOVA, *post hoc* Dunnett); error bars indicate standard deviation; n = 3. (D) CALS1 is required for NaCl-induced callose deposition. Comparisons are between untreated and NaCl-treated samples (one-way ANOVA, *post hoc* Tukey HSD); n = at least 6. (E) Impact of PLD on callose deposition in CRK2 lines. Comparisons are between untreated and NaCl-treated samples, pre-treated with 1-butanol or 2-butanol, for each line (one-way ANOVA, *post hoc* Tukey HSD); n = at least 6. (F) CRK2 can phosphorylate the N terminal of CALS1 *in vitro*, but cannot phosphorylate PLDα1. (G) Plasmodesmal permeability during standard growth conditions; the observed callose deposition correlates with changes in plasmodesmal permeability. Quantification by percent diffusion of a fluorescent intracellular dye from the adaxial to abaxial surface; comparisons are to Col-0 (one-way ANOVA, *post hoc* Dunnett); n = 3. Seven-day-old seedlings; NaCl treatment 150 mM 30 min; ns not significant, * P < 0.05, ** P < 0.01, *** P < 0.001.

The effect of CRK2 on salt-induced callose deposition was associated with active PLD-based PA production. No significant difference was found between the *pldα1* line and 1-butanol-treated Col-0, justifying the interpretation that 1-butanol treatment effectively inhibits PA production by PLD in this assay (Fig. 4E). PLD inhibition did not affect basal callose levels in any of the lines (Fig. 4B and E), but effectively prevented increases in callose deposition following NaCl treatment in all lines (Fig. 4E). Again, 2-butanol was used as a negative control as it does not inhibit PLD activity. A callose response similar to untreated conditions was observed following 2-butanol pre-treatment (Fig. 4E). Therefore, PLD, like CRK2, is required for the salt-induced callose response. Since CRK2 kinase activity was required for the salt-induced callose response, we tested the direct phosphorylation capability of CRK2 *in vitro*, and found that CRK2 could phosphorylate CALS1 but not PLDα1 (Fig. 4F). This suggests phosphorylation as a means by which CRK2 might regulate CALS1, and points to CRK2 downstream of PLDα1 and upstream of CALS1 as the mostly likely signalling cascade.

To test whether the observed callose deposition had a biological effect on plasmodesmal permeability, a modified version of the Drop-And-See (DANS) assay was employed (Lee et al., 2011). In this assay a fluorescent intracellular dye is applied to the adaxial surface of the leaf and diffusion to the abaxial surface is assessed, representing the relative permeability of plasmodesmata. Overexpression of CRK2 resulted in increased plasmodesmal permeability under standard growth conditions (Fig. 4G), in accordance with the decreased basal callose deposition observed in this line (Fig. 4A and B). The *crk2* line showed an intermediate phenotype where plasmodesmal permeability did not significantly differ from Col-0, perhaps due to genetic redundancy (Fig. 4G). The *pldα1* and *cals1* lines did not significantly differ from Col-0 under standard growth conditions (Fig. 4G), suggesting their role at plasmodesmata may be relevant primarily during salt stress.

### CRK2 is not required for the initial salt-induced calcium response

A common feature between CRK2 re-localization, callose deposition, and PLD activation is the requirement for Ca^2+^. Therefore, we investigated if CRK2 affected calcium signalling directly using calcium imaging. Fluo-4-AM (Molecular Probes; Thermo Fisher Scientific, Waltham, USA), a calcium-sensitive fluorescent probe which can be transiently loaded into cells, was used to measure the salt-induced calcium response in epidermal cells. The calcium response to NaCl treatment appeared highly similar in both Col-0 and *crk2* (Fig. 5A), suggesting that CRK2 is not likely to play a crucial role in regulation of the initial calcium elevation during salt stress. Interestingly, *crk2* exhibited an increased response to mock treatment when compared to Col-0 (Fig. 5A). This might suggest a higher mechanosensitivity in *crk2*. One drawback of using transient probes is the potential for differential probe loading between samples and genotypes. Therefore we also used stable transgenic plant lines expressing the FRET (Förster resonance energy transfer)-based calcium sensor yellow chameleon YCNano-65 (Horikawa et al., 2010). Calcium imaging was performed on the adaxial epidermal tissue layer. Again, no strong differences were observed between YCNano-65/Col-0 and YCNano-65/*crk2* during the initial calcium response to NaCl treatment (Fig. 5B). Together, these results support the hypothesis that CRK2 is not directly affecting the initial calcium signal itself during salt stress, but is acting downstream in a calcium-dependent manner.

**Fig. 5.**
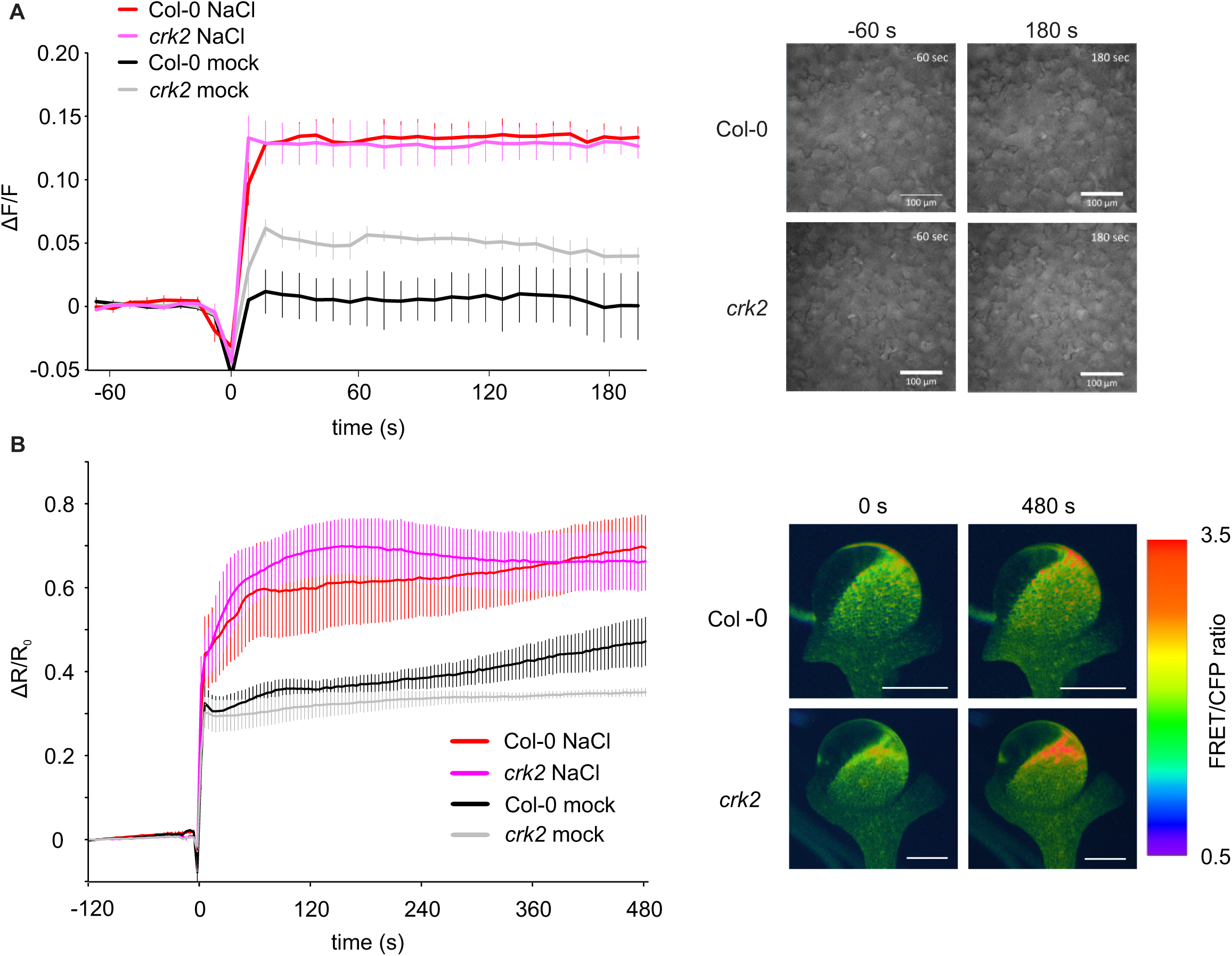
CRK2 is not required for the initial salt-induced calcium response. (A) Fluo-4-AM calcium imaging of cell-level Ca^2+^ influx in response to 10 mM NaCl; seven-day-old seedlings, epidermal cells; n = 3, approximately 70 cells measured per replicate. Scale bar = 100 µm. (B) YCNano-65 calcium imaging of tissue-level Ca^2+^ influx in response to 150 mM NaCl; seven-day-old seedlings; n = at least 6. Scale bar = 1 mm. Additions were made at t = 0. Error bars indicate standard error of the mean.

## Discussion

Soil affected by high salinity is becoming progressively more widespread, particularly across irrigated agricultural land. This poses an increasing threat to agricultural productivity, as the majority of crop species are not inherently salt tolerant (Yang and Guo, 2017). However, attempts at increasing salt tolerance, through breeding or genetic engineering, must first be preceded by a more thorough understanding of the mechanisms underlying salt tolerance and the molecular pathways involved in salt sensing and cellular responses. As discussed earlier, RLKs are responsible for much of the communication between the extracellular and intracellular environment, and the CRKs specifically have been implicated in various stress responses (Bourdais et al., 2015). Based on phylogenetic analysis, the CRKs can be split into two major groups: basal CRKs and variable CRKs. CRK2 is a member of the basal group of CRKs, which show considerable evolutionary conservation across plant species (Vaattovaara et al., 2019). Large scale phenotyping of the *crk2* mutant revealed changes in development, biotic stress responses, and abiotic stress responses, including high salinity (Bourdais et al., 2015). Building on this knowledge, the next step is to characterize the specific functions and protein interactions of CRK2 in these processes.

Callose deposition is central to the regulation of plasmodesmal permeability and therefore cell-to-cell communication. This is crucial not only for normal plant development and cellular signalling, but also during adverse conditions where the plant may choose to either close off communication to isolate an affected cell, or open communication to allow distant cells to respond accordingly. Increased callose deposition has been documented for various stresses (Wu et al., 2018), but has not yet been clearly linked to salt stress responses. Here we show that callose deposition is elevated during salt tolerance and the acute response to salt stress, and that NaCl-induced callose deposition is regulated in part by CRK2. Increased CRK2 expression leads to higher levels of callose deposition following salt treatment, and functional, kinase-active CRK2 protein is required for the salt-induced callose response. The callose synthases interacting with CRK2 (CALS1, CALS3, and CALS12) all contain at least one predicted phosphorylation site (PhosPhAT 4.0, http://phosphat.uni-hohenheim.de/; Durek et al., 2010; Heazlewood et al., 2008; Zulawski et al., 2013), and CRK2 can phosphorylate CALS1 *in vitro*. Thus, phosphorylation could be an important means to regulate callose synthase activity, and an interesting subject for future research. PLD activity was also required for salt-induced callose deposition; however, it did not affect the basal callose levels. This suggests that PLD may not be directly involved in the CRK2–callose synthase interaction, but is more likely exerting its effect upstream in this pathway.

The PLD protein family has been previously associated with salt stress tolerance (Hong et al., 2010). These enzymes cleave phospholipids to produce PA and a free head group. PLD, along with its product PA, has been linked to a wide range of cellular processes in eukaryotic cells, including endocytosis and vesicle trafficking (Koch et al., 2003; Lee et al., 2006; Shen et al., 2001; Thakur et al., 2016), membrane composition and microdomains (Faraudo and Travesset, 2007), and microtubule and cytoskeletal dynamics (Zhang et al., 2017, 2012). Twelve PLD genes are present in Arabidopsis, of which PLDα1 is the major isoform (Pappan et al., 1998, 1997a, 1997b; Qin and Wang, 2002; Qin et al., 1997; Wang and Wang, 2001). Salt stress induces the expression of multiple PLD genes (Katagiri et al., 2001), and the *pldα1, pldα3*, and *pld*δ mutants have been previously characterized as salt sensitive (Bargmann et al., 2009; Hong et al., 2008; Yu et al., 2010). Increased PA concentrations have been documented following various abiotic and biotic stresses, including hyperosmotic, salt, drought, freezing, wounding, and pathogens (Testerink and Munnik, 2005). However, in many cases the downstream targets and effectors of PLD/PA signalling are unknown in these responses. PLDα1 was consistently found as one of the top interacting proteins with CRK2 and many of CRK2’s cellular functions are dependent on PLD activity, as are the subcellular localization changes. Arabidopsis lines lacking functional PLDα1 or CRK2 have similarly decreased salt tolerance during germination, and several of the CRK2-affected cellular phenotypes have also been linked to PLDα1. The inhibitor results and the observation that CRK2 cannot phosphorylate PLDα1 support the hypothesis that it is PLD activity, and PA production, which is a key factor in the interaction of CRK2 and PLD, rather than a CRK2-directed phosphorylation-based interaction.

PLD activity can influence microtubules and the cytoskeletal structure as part of the response to biotic and abiotic stress. Activation of PLDα1 triggers microtubule depolymerization during abscisic acid (ABA)-induced stomatal closure (Jiang et al., 2014) and stabilizes microtubule organization during salt stress (Zhang et al., 2012). In plant cells, the cytoskeleton extends through plasmodesmata to neighboring cells. Modulation of cytoskeletal components actin and myosin at the neck regions of plasmodesmata may play a role in regulating plasmodesmal permeability, in addition to callose deposition (White and Barton, 2011). It was also shown that different types of plasmodesmata do not all respond the same to actin and/or myosin inhibitors, and thus the cytoskeleton may also be important for regulating plasmodesmal transport specificity and localizing molecules to plasmodesmata (White and Barton, 2011). PA itself influences membrane properties due to its negative charge, binding capacity for divalent cations (Faraudo and Travesset, 2007), and ability to induce membrane curvature (Kooijman et al., 2003). This curvature is important for example during exocytosis and endocytosis, where high degrees of curvature are required for membrane budding and vesicle formation, as well as protein localization (Zhao et al., 2017). PA-rich membrane domains can also directly act as localization signals for PA binding proteins, such as the sphingosine kinase 1 (Delon et al., 2004). Specialized microdomains at plasmodesmata have already been described, and appear essential for targeting of many plasmodesmata-localized proteins (Nicolas et al., 2018). Thus, PLD, through its action on membrane domains and cytoskeletal dynamics, could provide a mechanism for the CRK2 localization changes, and for bringing together various components such as CRK2 and CALS1.

In our proposed model, increased extracellular NaCl concentrations would trigger Ca^2+^ influx, as well as ROS production, as one of the earliest initial responses. This Ca^2+^ signal activates PLDα1 and causes its translocation from the cytoplasm to plasma membrane, leading to PA production and a shift in membrane properties (Fig. 6). This would serve as the scaffold for a change in CRK2 localization from uniformly along the plasma membrane to specific domains concentrated at plasmodesmata. Once localized at plasmodesmata, CRK2 interacts with CALS1 to promote callose deposition, ultimately leading to enhanced salt tolerance (Fig. 6). This could explain the observation that NaCl-induced callose deposition – and the variation between the differentially expressed CRK2 lines – is dependent on active PLD, but basal callose deposition is not affected by PLD activity. It also serves to link CRK2 protein function with the dynamic subcellular localization observed.

**Fig. 6.**
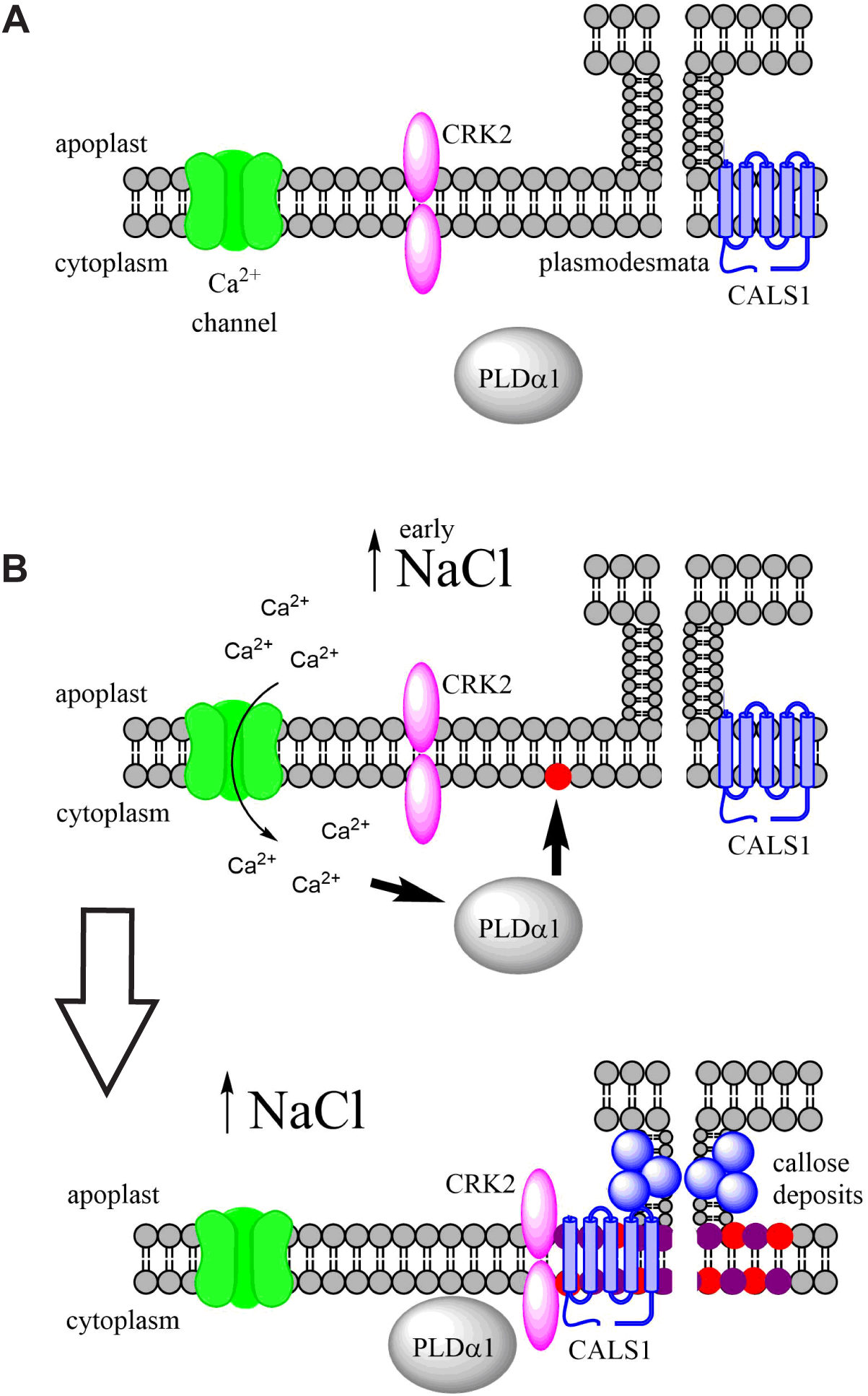
Schematic of proposed pathway for CRK2 regulation of callose deposition at plasmodesmata during salt stress. (A) Resting state. (B) Early responses to salt stress. Increased extracellular NaCl triggers Ca^2+^ influx. Cytoplasmic Ca^2+^ elevation activates PLDα1 leading to PA production and a shift in membrane properties; this serves as a scaffold for changes in CRK2 localization from uniformly along the plasma membrane to specific domains concentrated at plasmodesmata. Once localized at plasmodesmata, CRK2 interacts with CALS1 to promote callose deposition, ultimately leading to enhanced salt tolerance.

Here we have shown that CRK2 enhances salt tolerance at the germination stage in *Arabidopsis thaliana* and also increases root length under salt conditions. We demonstrated that CRK2 is involved in the regulation of callose deposition and can interact with CALS1. We found significant differences in callose deposition between wild type, *crk2* mutant, and CRK2 overexpressing lines, which correlate with differences in plasmodesmal permeability, and established that functional CRK2 is required for salt-induced callose deposition. In doing so, we revealed a novel role for callose deposition, in response to increased salinity, and demonstrated its importance for salt tolerance during germination. Using fluorescently tagged proteins we observed specific changes in CRK2’s subcellular localization in response to various stress treatments. These functions and localization are dependent on CRK2 kinase activity, as well as calcium and active PLD-based PA production. Thus we propose that CRK2 acts downstream of PLDα1 during salt stress to promote callose deposition and regulate plasmodesmal permeability, and that CRK2 adopts specific stress-dependent subcellular localization patterns in order to carry out its functions.

## Methods

### Growth Conditions

For all experiments, seeds were surface sterilized and plated under sterile conditions on half strength Murashige and Skoog media (Sigma-Aldrich, Darmstadt, Germany) supplemented with 0.8% agar, 1% sucrose, and 0.1% MES, pH 5.8. For selection of transgenic lines 20 µg/mL Basta (DL-phosphinothricin; Duchefa Biochemie, Haarlem, The Netherlands) and 100 µg/mL ampicillin was added. Plants were grown in a Sanyo growth chamber (Osaka, Japan) with a 16 h light 8 h dark photoperiod. For the DANS assay, seedlings were transferred to soil (2:1 peat:vermiculite) in the greenhouse after 7 days. For seed propagation and transgenic line creation, seeds were germinated on soil in the greenhouse and grown with a 12 h light 12 h dark photoperiod. All seeds were stratified in darkness at 4°C for at least 2 days.

### Plant Lines and Constructs

Col-0 was used as wild type for all experiments. T-DNA insertion lines for *crk2* (SALK_012659C; At1g70520), *pldα1* (SALK_053785; At3g15730), and *cals1-5* (SAIL_1_H10; At1g05570) were obtained from NASC European Arabidopsis Stock Centre (University of Nottingham, United Kingdom). Additional *cals1-2* and *cals1-3* alleles were obtained from Cui & Lee, 2016. Constructs for CRK2–YFP and YFP–Myc fusion proteins were created using the MultiSite Gateway technology (Invitrogen; Thermo Fisher Scientific, Waltham, USA). The coding sequence of CRK2 was amplified by PCR (forward primer: ATGAAGAAAGAACCTGTCC, reverse primer: TCTACCATAAAAGGAACTTTGTGAG) and inserted into pDONRzeo (Invitrogen; Thermo Fisher Scientific), then transferred to the pBm43GW (Invitrogen; Thermo Fisher Scientific) expression vector. The promoter region of CRK2 was amplified by PCR (forward primer: GGTTTTAGATCGTGTTAGATATATCA, reverse primer: TTTGTTTTGTTTGATTGAGAAA) and inserted into pDONR4R1 (Invitrogen; Thermo Fisher Scientific), then transferred to pBm43GW. mVenusYFP, RFP, Myc, PDLP5, and the CaMV 35S promoter were transferred from existing donor vectors into pBm43GW. To create the kinase-dead protein variants, point mutations were introduced into the coding sequence using mutagenic primers on the donor vectors, and then transferred into pBm43GW. Transgenic lines were created via Agrobacterium-mediated floral dipping using *Agrobacterium tumefaciens* GV3101_pSoup. CRK2 and YFP overexpression lines were created in Col-0 background and CRK2 complementation and kinase-dead lines were created in the *crk2* background. Transformed seeds were selected by Basta resistance until T3 homozygous lines were obtained. Transgenic lines are identified as follows: 35S::CRK2–YFP_9-3/Col-0, 35S::YFP–Myc_9-6/Col-0, pCRK2::CRK2–YFP_1-22/*crk2*, pCRK2::CRK2–YFP_1-17/*crk2*, 35S::CRK2^K353E^–YFP_4-5/*crk2*, 35S::CRK2^D450N^–YFP_11-2/*crk2*.

### Genotyping and RT-PCR

Genomic DNA was extracted from seven-day-old seedlings and used as a template for PCR-based genotyping. Extraction buffer: 100 mM Tris-HCl pH 8.0, 50 mM EDTA, 500 mM NaCl. For RT-PCR, RNA was extracted from seven-day-old seedlings, followed by cDNA synthesis, as described previously (Bourdais et al., 2015); this cDNA was used as a template for the PCR reactions. PP2AA3 (At1g13320) was used as a reference gene. Primers used for genotyping and RT-PCR are as follows: CRK2 (GCTAACTATGGTCTTGCGCAG, CAAAGATGAATCGATCAAGGC), PLDα1 (CAAGGCTGCAAAGTTTCTCTG, CATCAATGCCCTGCACTTAAT), CALS1 (TTAGACATTCAGGGGTTCGTG, GACGAAAACATTGGTTCTCCA), PP2AA3 (GAGGATGTCTATGGTTGATG, GCCATTCCCATTATAACTG).

### Transient Expression in *Nicotiana benthamiana*

The pFLS2::FLS2–GFP construct was transformed into GV3101_pSoup Agrobacterium and infiltrated into the leaves of 6 week old *Nicotiana benthamiana* plants. The C58C1 Agrobacteria strain, containing P19, was co-infiltrated at a 1:1 ratio to enhance and prolong expression. Infiltration media: 10 mM MES pH 5.6, 10 mM MgCl_2_, 200 µM acetosyringone. The maximum expression was observed at 2 days post-infiltration, and was the time point used for all experiments. Leaf discs were cut from infiltrated areas and transferred to 12-well plates for treatments (Table 2).

### Transient Transformation of Arabidopsis Seedlings

The constructs 35S::CRK2–YFP_pBm43GW and 35S::PDLP5–RFP_pBm43GW were transformed into GV3101_pSoup Agrobacteria and then transiently transformed into Arabidopsis seedlings using the FAST co-cultivation method (Li et al., 2009). Following transformation, the seedlings were kept in darkness for 40 h, moved to light for 24 h, and then analyzed by microscopy as seven-day-old seedlings.

### Immunoprecipitation and Mass Spectrometry

Immunoprecipitation experiments were performed as described previously (De Rybel et al., 2013; Zwiewka et al., 2011), using 0.5 g of seven-day-old seedlings, collected under normal conditions, NaCl, or H_2_O_2_ treatments (Table 2). Interacting proteins were isolated from total protein extracts using anti-GFP-coupled magnetic beads (Miltenyi Biotec, Bergisch Gladbach, Germany). Proteins were digested with trypsin to peptides, purified, and sent for identification by mass spectrometry (MS). The MS analyses were performed on a nanoflow HPLC system (Easy-nLC1000, Thermo Fisher Scientific) coupled to the Q Exactive mass spectrometer (Thermo Fisher Scientific, Bremen, Germany). Peptides were first loaded on a trapping column and subsequently separated inline on a 15 cm C18 column (75 μm x 15 cm, ReproSil-Pur 5 μm 200 Å C18-AQ). The mobile phase consisted of water with 0.1% formic acid (solvent A) or acetonitrile/water (80:20 (v/v)) with 0.1% formic acid (solvent B). A 50 min gradient from 6% to 43% B was used to elute peptides. A constant 300 nl/min flow rate was used. MS data was acquired automatically by using Thermo Xcalibur 4.0 software (Thermo Fisher Scientific). A data dependent acquisition method consisted of an Orbitrap MS survey scan of mass range 300-1800 m/z, followed by HCD fragmentation for the 10 most intense peptide ions. Data files were searched for protein identification using Proteome Discoverer 2.1 software (Thermo Fisher Scientific) connected to an in-house server running the Mascot 2.5.1 search engine (Matrix Science, London, UK). The data was searched against the TAIR10 database. The 35S::CRK2–YFP_9-3 overexpression line was used in all replicates. Unspecific interactors were removed by comparison to the 35S::YFP–Myc_9-6 control line to exclude proteins identified as interacting with YFP. Only proteins with more than one peptide were considered as true identifications.

### Germination Assay

Seeds were germinated on either untreated, 150 mM NaCl, or 200 mM NaCl media and assessed on day 6. Percent germination was calculated by counting the number of germinated seeds versus total seeds. The untreated samples were set as 100% germination to allow for normalization of data and comparison between lines. Statistical significance was determined by one-way ANOVA with *post hoc* Dunnett’s test using JMP Pro 13 (SAS Institute Inc., Cary, USA). Three replicates were performed for each line and treatment.

### Root Length Assay

Seeds were germinated on regular growth media and transplanted on day five to either untreated or 150 mM NaCl media. Plates were grown in a vertical position and primary root length was measured on day eight. Statistical significance was determined by one-way ANOVA with *post hoc* Dunnett’s test using JMP Pro 13. Replicates are as indicated in the figure legends.

### Western Blot

Following treatments, plant material was immediately frozen in liquid nitrogen and ground to a fine powder. Total proteins were extracted with sodium dodecyl sulfate (SDS) extraction buffer (50 mM Tris-HCl pH 7.5, 2% SDS, 1% protease inhibitor cocktail (Sigma-Aldrich)) and centrifuged at 4°C, 16 000 x g for 20 min. Supernatants were loaded in equal protein concentrations and resolved by SDS-PAGE, then transferred to Immobilon-FL polyvinylidene difluoride (PVDF) membranes (Merck Millipore, Darmstadt, Germany). Western blotting was carried out using mouse anti-GFP 11814460001 (Roche, Basel, Switzerland) and rabbit anti-Histone H3 AS10710 (Agrisera, Vännäs, Sweden) primary antibodies, and goat anti-mouse IRDye800CW (LI-COR, Lincoln, USA) and goat anti-rabbit IRDye800CW (LI-COR) secondary antibodies, and imaged with the Odyssey Infrared Imaging System (LI-COR). Quantification of western blots was carried out in Image J (U. S. National Institutes of Health, Bethesda, USA; https://imagej.nih.gov/ij/) by measuring band mean intensity. Col-0 was set as the background level and protein levels were normalized to RUBISCO (Ribulose-1,5-bisphosphate carboxylase/oxygenase) and Histone H3 as internal controls.

### *In vitro* Kinase Assay

Constructs for 6His–GST–CRK2cyto (CRK2 cytoplasmic domain), 6His–MBP–PLDα1, and 6His–MBP–CALS1_N (CALS1 N-terminus) recombinant proteins were generated using In-Fusion technology (Clontech; Takara Bio USA, Mountain View, USA). The fragment of CRK2cyto (WT, K353E, or D450N) was amplified by PCR (forward primer: AAG TTCTGTTTCAGGGCCCGAAGAGGAAGAGAAGAGGATC, reverse primer: ATGGTCTAGAAAGCTTTATCTACCATAAAAGGAACTTTGTGA) from pDORNzeo-CRK2 plasmid and cloned into pOPINK vector. PLDα1 (forward primer: AAGTTCTGTTTCAGGGCCCGATGGCGCAGCATCTGTTGCACGGG, reverse primer: ATGGTCTAGAAAGCTTTATTAGGTTGTAAGGATTGGAGGCAGG) and CALS1_N (forward primer: AAGTTCTGTTTCAGGGCCCGATGGCTCAAAGAAGGGAACCTGATC, reverse primer: ATGGTCTAGAAAGCTTTATCTATCAAAACTTCTAAATATATGC) were amplified from cDNA and cloned into pOPINM vector. 6His–GST–CRK2cyto (WT, K353E and D450N) were expressed in *Escherichia coli* Lemo21 and purified by Glutathion Sepharose 4B (GE Healthcare, Chicago, USA). 6His–MBP–PLDα1 and 6His–MBP–CALS1_N were expressed in *E. coli* BL21 and purified by Amylose Resin (New England Biolabs, Ipswich, USA). One µg of kinase protein was incubated in kinase buffer (50 mM HEPES pH 7.4, 1 mM dithiothreitol, 10 mM MgCl_2_) for 30 min at room temperature with [γ-^32^P]-ATP and substrate protein. Myelin basic protein (Sigma-Aldrich) was used as an artificial substrate. The samples were subsequently separated by SDS-PAGE and exposed to an imaging plate overnight. Radioactivity scans were obtained with Fluor Imager FLA-5100 (Fujifilm, Tokyo, Japan).

### Subcellular Protein Localization

Stable homozygous lines expressing mVenusYFP-fusion proteins were used for all live imaging of CRK2. Transient expression of pFLS2::FLS2–GFP in *Nicotiana benthamiana* was used for FLS2 internalization controls. Transient transformation of Arabidopsis seedlings was used for CRK2 localization in the *pldα1* mutant background and for co-localization with PDLP5. Seven-day-old seedlings were transferred to 12-well plates and treatments applied as described in Table 2. Samples were mounted in the treatment solution and imaged immediately. Fluorescent images were obtained with Leica TCS SP5 II HCS confocal microscope using standard YFP settings (CRK2–YFP) of 514 nm excitation and a detection range of 525–590 nm, standard GFP settings (FLS2–GFP) of 488 nm excitation and a detection range of 500–600 nm, or standard RFP settings (PDLP5–RFP) of 561 nm excitation and a detection range of 560–600 nm. Quantification of CRK2 co-localization with callose deposits or PDLP5 was achieved using the following equation: % co-localization = (number of co-localized spots / total number of CRK2 spots). Quantification of CRK2–YFP re-localization was achieved by calculating the percent enrichment at the re-localization domains (with Image J) using the following equation: % enrichment = (fluorescent intensity “spot” / fluorescent intensity general plasma membrane) x 100. Statistical significance was determined by one-way ANOVA with pooled t-test test using JMP Pro 13. Replicates are as indicated in the figure legends.

### Callose Staining

Seven-day-old seedlings were transferred to 12 well plates for treatments (Table 2), then fixed overnight in 1:3 acetic acid:ethanol. Seedlings were washed with 150 mM K_2_HPO_4_ (Sigma-Aldrich) for 30 min and stained with 0.01% aniline blue (Sigma-Aldrich) + 150 mM K_2_HPO_4_ for 2 hours in darkness. Fluorescent images were obtained with Leica TCS SP5 II HCS confocal microscope using standard DAPI settings of 405 nm excitation and a detection range of 430–550 nm. The number of callose deposits was counted manually from each image area (780.49 µm^2^). Statistical significance was determined by one-way ANOVA with *post hoc* Tukey HSD using JMP Pro 13. Replicates are as indicated in the figure legends. Image intensity was enhanced for visual representation, however all quantifications were made from the original images.

### DANS Assay for Plasmodesmata Permeability

Experiments were performed using a modified version of the DANS assay (Lee et al., 2011). Briefly, rosette leaves were cut from three-week-old plants and a 1 µL drop of 5 µM fluorescein diacetate was applied to the adaxial surface. After 5 minutes the liquid was removed with filter paper, samples mounted in water, and imaged immediately. Fluorescent images were obtained with Leica TCS SP5 II HCS confocal microscope using standard GFP settings of 488 nm excitation and a detection range of 500–600 nm. Percent diffusion was calculated by dividing the average total fluorescence from abaxial images by the adaxial images. Statistical significance was determined by one-way ANOVA with *post hoc* Dunnett’s test using JMP Pro 13. Three replicates were performed for each line.

### Calcium Imaging

The fluo-4-AM (Molecular Probes; Thermo Fisher Scienctific) synthetic calcium probe was used for cell-level calcium imaging. Cotyledons were removed from seven-day-old seedlings and placed in 96-well plates. Cells were loaded for 1 h at +4°C in darkness, in loading buffer composed of 10 mM MES, 2 mM probenecid, and 5 µM fluo-4-AM mixed 1:1 with 20% (w/v) pluronic acid. The cotyledons were then washed with experimental buffer (10 mM MES, 2 mM probenecid) and mounted to an open microscope slide chamber, with a 3.0 µm pore size polycarbonate filter (Polycarbonate 3.0 micron; Osmonics, Minnetonka, USA) on top for immobilization (Shariatmadari et al., 2001). Epidermal cells from the adaxial surface were observed. Calcium imaging was performed with a Nikon TE2000 fluorescence microscope. Samples were exposed to 480 nm excitation, and the emitted light was collected through a 505 nm dichroic mirror and a 510–560 nm band pass filter. Images were acquired every 8 seconds and treatments were added directly to the chamber during imaging. A final concentration of 150 mM NaCl was used for treatments; mock treatments consisted of experimental buffer. The data were normalized using the following equation: ΔF/F_b_ = (F_t_ – F_b_)/F_b_, where F_t_ is the fluorescence measured at time point t and F_b_ is the baseline fluorescence. The baseline was taken as an average of the eight measurements prior to the addition. Approximately 70 cells were measured each run and experiments were repeated three times. Analyses and graphs were made with Microsoft Excel.

Plants expressing the genetically encoded YCNano-65 calcium probe were used for tissue level calcium imaging. YCNano-65/Col-0 was provided by Prof. Simon Gilroy (University of Wisconsin, USA) and has been previously described (Choi et al., 2014). YCNano-65/*crk2* was generated by crossing YCNano-65/Col-0 with the *crk2* T-DNA mutant. Homozygous F3 lines were selected by Basta resistance (YCNano-65 insertion) and genotyping (T-DNA insertion). Genotyping primers for *crk2* are described by (Bourdais et al., 2015). Seven-day-old seedlings were mounted and 1 µL of 150 mM NaCl or MS media (mock treatment) was applied to the adaxial surface of cotyledons. Calcium imaging was performed with a Nikon SMZ25 microscope using the settings described previously (Lenglet et al., 2017). CFP and FRET (cpVenus) images were acquired simultaneously every 4 seconds. Data is presented as the ratio of FRET to CFP signal and was normalized to the initial baseline using the following equation: ΔR/Ro = (R_t_ – Ro)/Ro, where R_t_ is the ratio value at time point t and Ro is the initial ratio value. Experiments were repeated at least six times. Analyses and graphs were made with Microsoft Excel.

Schematic figures were made with ChemDraw 16 (PerkinElmer, Waltham, USA).

## Author Contributions

KH designed experiments, performed experiments, analyzed data, and wrote the manuscript. SK, AR, CT, and MT performed experiments. JPK and MW designed experiments. All authors contributed to and edited the final manuscript.

## Acknowledgements

The authors would like to thank Drs. Alexey Shapiguzov and Julia Krasensky-Wrzaczek for critical comments on the manuscript. We thank Tuomas Puukko, Nghia Le Tri, and Jiaqi Wang (Saitama University, Japan) for technical assistance, Dr. Riccardo Siligato for the Gateway Multisite vector system, and Prof. Jung-Youn Lee (University of Delaware, USA) for the *cals1* mutant seeds. Microscopy imaging was performed at the Light Microscopy Unit, Institute of Biotechnology, University of Helsinki. Mass spectrometry analyses were performed at the Turku Proteomics Facility, supported by Biocenter Finland. This work was supported by the Academy of Finland (grant numbers #275632, #283139, and #312498 to MW), the University of Helsinki (Three-year fund allocation to MW), and KAKENHI (17H05007, 18H04775, and 18H05491 to MT). KH, SK and MW are members of the Centre of Excellence in the Molecular Biology of Primary Producers (2014-2019) funded by the Academy of Finland (grant numbers #271832 and #307335).

## Supplementary Table Legends

**Table S1. Full list of proteins identified as interacting with CRK2.** The 35S::CRK2–YFP_9-3 overexpression line was used in all replicates; unspecific interactors were removed by comparison to the 35S::YFP–Myc_9-6 control line to exclude proteins identified as interacting with YFP. Each replicate represents a separate immunoprecipitation. Proteins with only 1 peptide identified were removed from further analysis (grey text). Seven-day-old seedlings; nt = untreated, NaCl = 150 mM NaCl 30 min, H2O2 = 1 mM H_2_O_2_ 30 min.

**Table S2. Gene ontology classifications (Biological Process) for identified CRK2 interacting proteins.** The 35S::CRK2–YFP_9-3 overexpression line was used in all replicates; unspecific interactors were removed by comparison to the 35S::YFP–Myc_9-6 control line to exclude proteins identified as interacting with YFP; proteins with only 1 peptide identified were excluded. Replicates of each treatment were combined. Seven-day-old seedlings; nt = untreated, NaCl = 150 mM NaCl 30 min, H2O2 = 1 mM H_2_O_2_ 30 min.

**Table S3. Semi-quantification of CRK2**–**YFP localization changes.** Degree of re-localization classified as none (-), weak (+), strong (++), very strong (+++). 35S overexpression lines used in all replicates; seven-day-old seedlings, epidermal cells; treatment times and conditions according to Table 2.

