## Supplementary figures 1-9 for "CRK2 enhances salt tolerance by regulating callose deposition in connection with PLDα1"

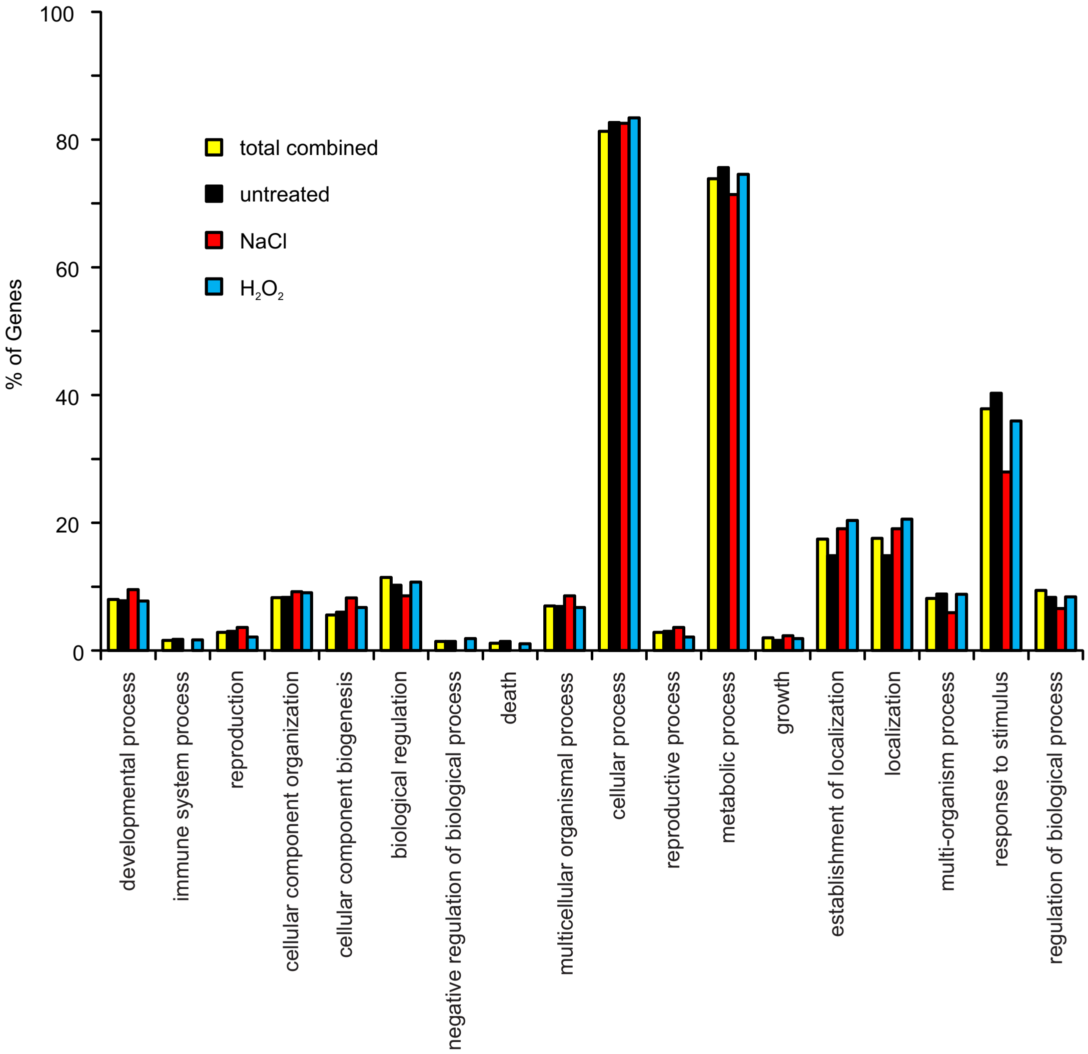


**Fig. S1. Gene ontology singular enrichment analysis of identified CRK2 interacting proteins.** Gene ontology categories based on biological process. 35S::CRK2-YFP_9-3 overexpression line used in all replicates; unspecific interactors were removed by comparison to the 35S::YFP-Myc_9-6 control line; proteins with only 1 peptide identified were excluded. Replicates of each treatment were combined. Seven-day-old seedlings; NaCl 150 mM 30 min, $H_2O_2$ 1 mM 30 min.

**A**

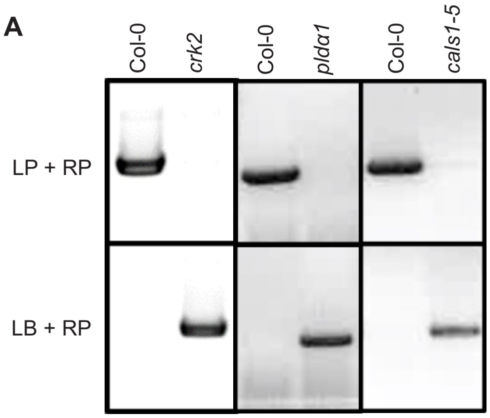


**B**

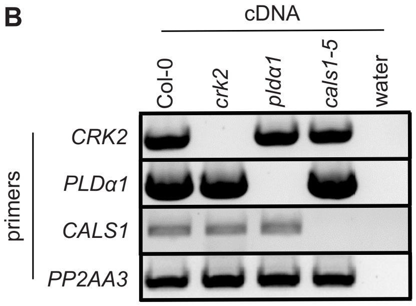


**Fig. S2. Confirmation of T-DNA mutant lines.** (A) Genotyping shows the presence of a T-DNA insert in the correct location. (B) RT-PCR shows absence of a transcript in the corresponding mutant, but presence in other mutants. PP2AA3 was used as a reference gene; the same cDNA template was used for each line with all primer sets.

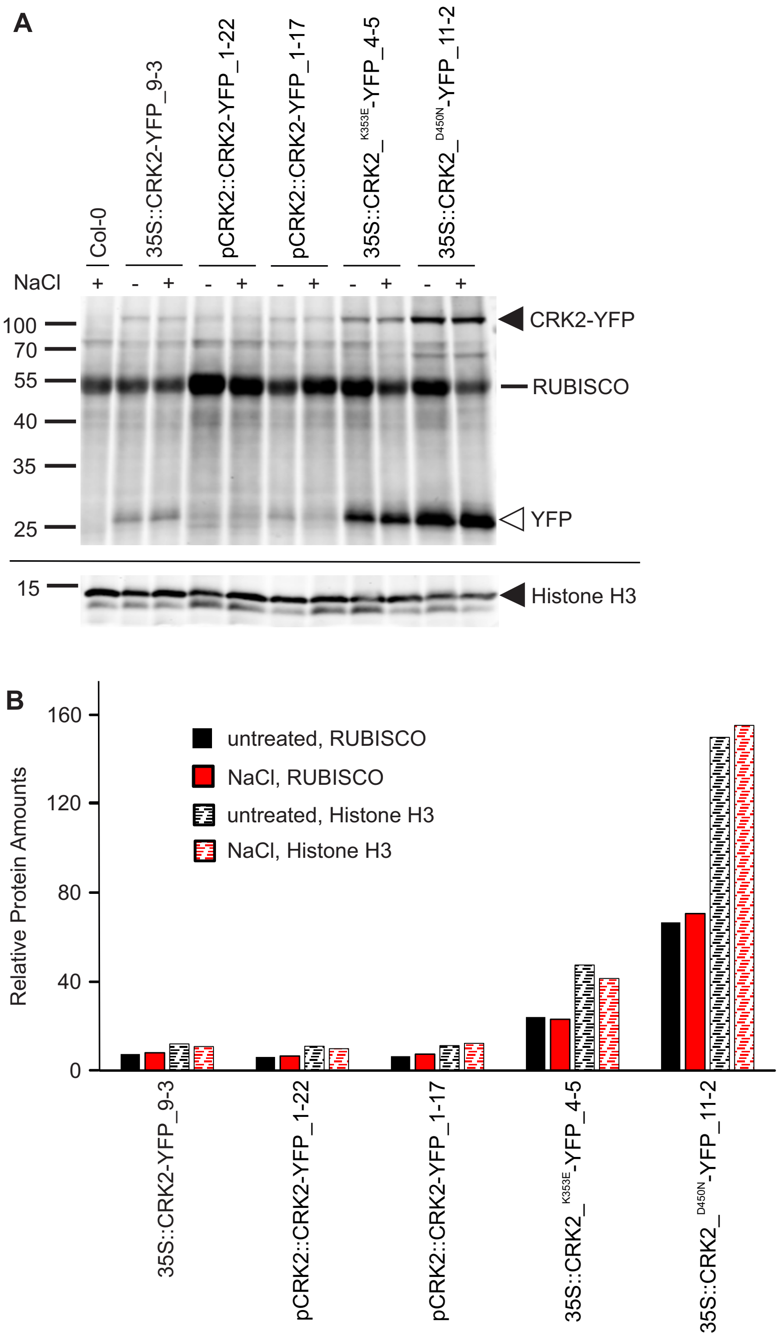


**Fig. S3. Protein expression in CRK2 transgenic lines.** (A) Western blot analysis. Upper blot shows the abundance of CRK2-YFP in transgenic plants. Lower blot shows the abundance of Histone H3 in the same extracts from transgenic plant lines. (B) Quantification of western blot; Col-0 was set as the background level and CRK2 bands were normalized to RUBISCO (solid bars) and Histone H3 (hatched bars) as internal controls.

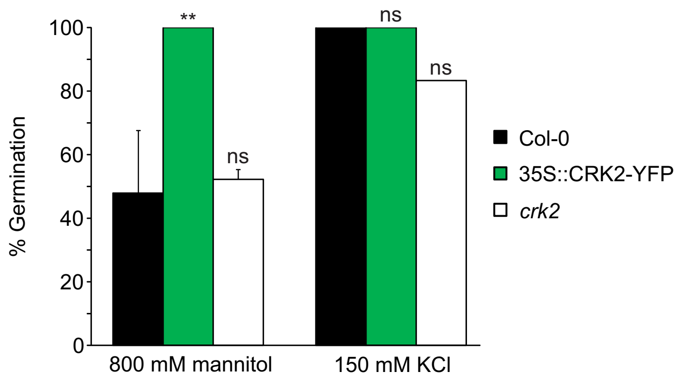


**Fig. S4. The osmotic component has a greater effect than the ionic component on the CRK2 mediated germination phenotype.** Data was normalized to the untreated controls for each line. Comparisons are to Col-0; ns not significant, * $P < 0.05$, ** $P < 0.01$, *** $P < 0.001$ (one-way ANOVA, *post hoc* Dunnett); n = 3; error bars indicate standard deviation.

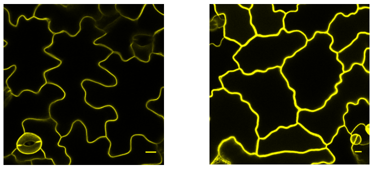

35S::CRK2$^{K353E}$-YFP 35S::CRK2$^{D450N}$-YFP

**Fig. S5. CRK2-YFP kinase-dead variants have proper subcellular localization.** 35S::CRK2$^{K353E}$-YFP and 35S::CRK2$^{D450N}$-YFP localize uniformly to the plasma membrane under standard growth conditions. Seven-day-old seedlings, epidermal cells. Scale bar = 10 μm.

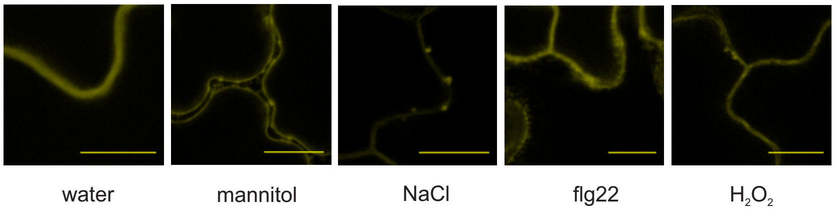


**Fig. S6. CRK2 expressed under its native promoter shows the same localization patterns as the overexpression line.** pCRK2::CRK2-YFP; seven-day-old seedlings, epidermal cells; mannitol 800 mM 15 min, NaCl 150 mM 30 min, flg22 10 μM 30 min, $H_2O_2$ 1 mM 30 min. Scale bar = 10 μm.

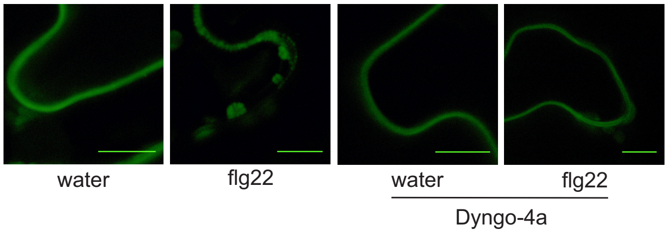


**Fig. S7. Dyngo-4a effectively inhibits FLS2 internalization following flg22 treatment.** pFLS2::FLS2GFP transiently expressed in *Nicotiana benthamiana;* epidermal cells; flg22 10 µM 30 min. Scale bar = 10 µm.

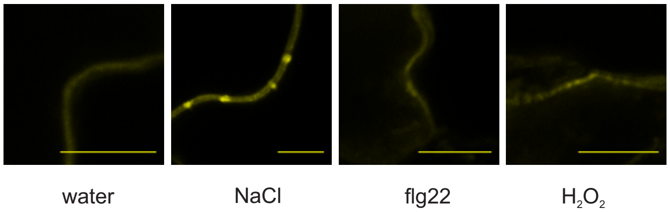


**Fig. S8. CRK2 expressed transiently shows the same localization patterns as the stable overexpression line.** 35S::CRK2-YFP transiently expressed in Col-0 via agrobacteria seedling co-cultivation; seven-day-old seedlings, epidermal cells; NaCl 150 mM 30 min, flg22 10 µM 30 min, $H_2O_2$ 1 mM 30 min. Scale bar = 10 µm.

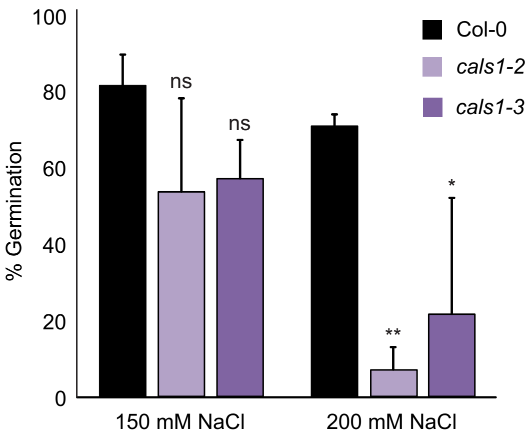


**Fig. S9. Germination response to salt of additional *cals1* alleles.** Data was normalized to the untreated controls for each line. Comparisons are to Col-0; ns not significant, * $P < 0.05$, ** $P < 0.01$, *** $P < 0.001$ (one-way ANOVA, *post hoc* Dunnett); n = 3; error bars indicate standard deviation.
