## Supplementary table 3 for "CRK2 enhances salt tolerance by regulating callose deposition in connection with PLDα1"

**Table S3. Semi-quantification of CRK2−YFP localization changes.** Degree of re-localization classified as none (-), weak (+), strong (++), very strong (+++). 35S overexpression lines used in all replicates; seven-day-old seedlings, epidermal cells; treatment times and conditions according to Table 2.

| **Pre-Treatment** | **Treatment** | **Degree of Re-Localization** | **Corresponding Figure** |
| --- | --- | --- | --- |
|  | water | - | 2B |
|  | mannitol | +++ | 2B |
|  | NaCl | ++ | 2B |
|  | flg22 | + | 2B |
|  | H_2_O_2_ | + | 2B |
| Kinase-dead (CRK2^K353E^) | water | - | 3A |
| Kinase-dead (CRK2^K353E^) | NaCl | - | 3A |
| Kinase-dead (CRK2^K353E^) | flg22 | - | 3A |
| Kinase-dead (CRK2^K353E^) | H_2_O_2_ | - | 3A |
| DPI | water | - | 3B |
| DPI | NaCl | ++ | 3B |
| DPI | flg22 | - | 3B |
| DPI | H_2_O_2_ | + | 3B |
| LaCl_3_ | water | - | 3C |
| LaCl_3_ | NaCl | - | 3C |
| LaCl_3_ | flg22 | - | 3C |
| LaCl_3_ | H_2_O_2_ | - | 3C |
| DXM | water | - | 3C |
| DXM | NaCl | - | 3C |
| DXM | flg22 | - | 3C |
| DXM | H_2_O_2_ | - | 3C |
| EGTA | water | - | 3C |
| EGTA | NaCl | ++ | 3C |
| EGTA | flg22 | - | 3C |
| EGTA | H_2_O_2_ | - | 3C |
|  | DMSO | - | 3D i |
|  | CaCl_2_ | - | 3D ii |
|  | CaCl_2_ + ionomycin | ++ | 3D iii |
|  | CPA | + | 3D iv |
| DPI | CaCl_2_ + ionomycin | + | 3D v |
| DPI | CPA | + | 3D vi |
| Kinase-dead (CRK2^K353E^) | CaCl_2_ + ionomycin | ++ | 3D vii |
| Dyngo-4a | water | - | 3E |
| Dyngo-4a | NaCl | + | 3E |
| Dyngo-4a | flg22 | + | 3E |
| Dyngo-4a | H_2_O_2_ | + | 3E |
| 1-butanol | water | - | 3F |
| 1-butanol | NaCl | - | 3F |
| 1-butanol | flg22 | - | 3F |
| 1-butanol | H_2_O_2_ | - | 3F |
| 2-butanol | water | - | 3G |
| 2-butanol | NaCl | ++ | 3G |
| 2-butanol | flg22 | + | 3G |
| 2-butanol | H_2_O_2_ | + | 3G |
| *pldɑ1* background | water | - | 3H |
| *pldɑ1* background | NaCl | - | 3H |
| *pldɑ1* background | flg22 | - | 3H |
| *pldɑ1* background | H_2_O_2_ | - | 3H |
